# Combinatorial Tissue Engineering Partially Restores Function after Spinal Cord Injury

**DOI:** 10.1101/254821

**Authors:** Jeffrey S. Hakim, Brian R. Rodysill, Bingkun K. Chen, Ann M. Schmeichel, Michael J. Yaszemski, Anthony J. Windebank, Nicolas N. Madigan

## Abstract

Hydrogel scaffolds provide a beneficial microenvironment in transected rat spinal cord. A combinatorial biomaterials based strategy provided a microenvironment that facilitated regeneration while reducing foreign body reaction to the 3-dimensional spinal cord construct. We used poly lactic-co-glycolic acid microspheres to provide sustained release of rapamycin from Schwann cell (SC)-loaded, positively charged oligo-polyethylene glycol fumarate scaffolds. Three dose formulations of rapamycin were compared to controls in 53 rats. We observed a dose-dependent reduction in the fibrotic reaction to the scaffold and improved functional recovery over 6 weeks. Recovery was replicated in a second cohort of 28 animals that included retransection injury. Immunohistochemical and stereological analysis demonstrated that blood vessel number, surface area, vessel diameter, basement membrane collagen, and microvessel phenotype within the regenerated tissue was dependent on the presence of SCs and rapamycin. TRITC-dextran injection demonstrated enhanced perfusion into scaffold channels. Rapamycin also increased the number of descending regenerated axons, as assessed by Fast Blue retrograde axonal tracing. These results demonstrate that normalization of the neovasculature was associated with enhanced axonal regeneration and improved function after spinal cord transection.

## Introduction

Spinal cord injury (SCI) is a devastating condition affecting between 250,000 and 500,000 new people each year in the developed world (2013). The poor regenerative capacity of the human central nervous system is built on maintaining stability that is a biological advantage for a complex nervous system built on billions of interneuronal connections established during growth and development. This contrasts with the simpler peripheral nervous system that effectively regenerates after many types of injury. Failure of repair following SCI results from many factors. Glial and stromal scarring formed at the lesion site block axon growth and there is an increase in inhibitors associated with myelin debris and proteoglycan deposition in the lesion environment (Filbin 2003; Silver and Miller 2004). Acute trauma is followed by a local inflammatory response that causes additional tissue damage. This inflammatory and fibrotic response effectively walls off the injured area and protects surrounding surviving spinal tissue.

The intrinsic capacity for central axons to regenerate was demonstrated by Aguayo and colleagues more than 35 years ago (Richardson *et al*. 1980). We have developed a rationally designed combinatorial system to harness this intrinsic regenerative capacity. A biodegradable hydrogel scaffold and Schwann cells provide key elements of architecture and cellular support for a regenerative environment. We previously demonstrated that positively charged oligo[poly(ethylene glycol) fumarate] (OPF +) scaffolds loaded with or without Schwann cells facilitated a regeneration permissive lesion environment (Hakim *et al*. 2015). Barriers to axonal regeneration including the total area of stromal scarring, cyst formation, astrocyte reactivity, chondroitin sulfate proteoglycan (CSPG) deposition and myelin debris were all significantly reduced in rat spinal cords implanted with OPF+ scaffolds compared to animals with transection injury alone. We demonstrated significant axonal regeneration but not restoration of function. Over time scaffold channels became progressively occluded with collagen and fibrinoid scarring and the implanted Schwann cells migrated out of the scaffolds into the adjacent spinal cord. This demonstrated a need to reduce the foreign body reaction to implanted scaffolds as a step towards improving functional recovery. This foreign body response has been observed with many natural (Gao *et al*. 2013; Gros *et al*. 2010; Horn *et al*. 2007; Jain *et al*. 2006; King *et al*. 2010) and synthetic (Bakshi *et al*. 2004; Gautier *et al*. 1998; Hejcl *et al*. 2009; Tsai *et al*. 2004; Wong *et al*. 2008; Xu *et al*. 1999) materials. Strategies to reduce scarring such as methylprednisolone (Chen *et al*. 1996; Chvatal *et al*. 2008) and chondroitinase ABC to degrade CSPGs (Fouad *et al*. 2005; Hwang *et al*. 2011; Morice *et al*. 2002) have had limited success.

An important novel finding in this scaffold model supported an association between vascular distribution and axonal regeneration. Using unbiased stereology to provide physiological estimates of blood vessel morphology and distribution, we demonstrated that the formation of smaller, densely packed vessels which resemble the capillary structure in normal spinal cord was significantly associated with increased axonal regeneration.(Madigan *et al*. 2014). We hypothesized that a strategy to reduce the foreign body reaction and normalize capillary distribution in the regenerating tissue would lead to functional recovery.

Rapamycin is a potent immunosuppressive drug widely used to prevent allograft organ transplant rejection. It has also been used experimentally to reduce fibrosis in reaction to various implanted synthetic materials (Morice *et al*. 2002; Moses *et al*. 2003). Rapamycin has also been shown to reduce microglial activation, astrocyte reactivity, macrophage/neutrophil infiltration and TNF-α secretion in the SCI lesion environment (Goldshmit *et al*. 2015; Tateda *et al*. 2017). It induces autophagy and activates the Wnt/β-catenin signaling pathway while preserving neurons and promoting motor functional recovery after SCI (Gao *et al*. 2015). Rapamycin may also have a role in normalizing blood vessel distribution as has been proposed in cancer therapy. We are building on rational, combinatorial, tissue engineering strategies by delivering rapamycin from microspheres to reduce the foreign body reaction and enhance normal vascularization of regenerating tissue (Guba *et al*. 2002; Jain 2001; Jain 2005).

Scaffold repair in the spinal cord transection model allows manipulation and study of individual components of the microarchitecture and microenvironment in the regenerating spinal cord. Although it does not mimic acute blunt trauma to the spinal cord, it provides a system where the regeneration process can be deconstructed and the effect of varying individual components of the regeneration environment can be systematically studied.

## Materials and Methods

### Dose finding to determine an effective dose of rapamycin that inhibits fibroblast proliferation *in vitro*

We first determined the dose of rapamycin required to inhibit fibroblast proliferation *in vitro*. 30,000 normal rat kidney (NRK) 49-F fibroblast cells (ATTC, CRL-1570) per well were plated in 24 well tissue culture plates in Dulbecco’s modified essential media (DMEM) with 5% fetal bovine serum (FBS) and incubated at 37° C in a 5% CO_2_ humidified atmosphere. The following day, the cells were serum-starved by changing the media to DMEM with 0.1% FBS for 24 hours. Fibroblasts were stimulated to proliferate by treatment with 5 ng/mL recombinant human transforming growth factor β1 (TGFβ1) (R&D Systems, Minneapolis, Minnesota, USA) in DMEM with 0.1% FBS. Control cells were maintained in low serum media only. Vehicle control, 1 nM, 10 nM, 20 nM, 40 nM, 80 nM, 100 nM and 1 mM doses of rapamycin (Toronto Research Chemicals, Toronto, Ontario, Canada) were prepared in triplicate wells. After two days in culture, fibroblast proliferation was measured using the CellTiter 96 AQueous One Solution Cell Proliferation MTS Assay kit (Promega, Madison, Wisconsin, USA) according to the manufacturer’s instructions. Briefly, the media was removed from the cells and replaced with 400 μL DMEM with 0.1% FBS and 100 μL of [3-(4,5-dimethylthiazol-2-yl)-5-(3-carboxymethoxyphenyl)-2-(4-sulfophenyl)-2H-tetrazolium, inner salt] (MTS) and phenazine ethosulfate (PES) solution. After 2 hours 200 μL of supernatant was placed in a 96 well plate and absorbance at 490 nm was measured using a Spectra Max Plus 384 plate reader (Molecular Devices, Sunnyvale, California, USA). Absorbances at 490 nm were averaged for each group and expressed as a percentage of the negative control group.

### Effect of rapamycin on axonal growth *in vitro*

We then determined whether doses of rapamycin that inhibited fibroblast proliferation were permissive for axonal growth *in vitro* using a previously validated and well-established assay (Gill *et al*. 1998; Staff *et al*. 2013). Dorsal root ganglia (DRG) explants were isolated from embryonic day 15 Sprague Dawley rat pups (Harlan Laboratories, Indianapolis, Indiana, USA) as previously described. Four DRG explants were placed in collagen-coated 35 mm dishes in enhanced minimally-enriched media (EMEM) containing 10% FBS, 7 mg/mL glucose, 1.2 mM l-glutamine, and 8 ng/mL nerve growth factor (NGF) (R&D Systems) in triplicate per group. Explants were treated with rapamycin concentrations ranging from 1 nM to 100 nM or an equivalent volume of absolute ethanol vehicle. DRG neurite outgrowth was then assessed at 24 and 48 hours after plating as previously described (Conti *et al*. 2004). Phase contrast images of live DRG explants were taken to measure the longest neurite arc in each explant using NIH ImageJ analysis software and averaged for each treatment group for statistical analysis.

### Measurement of rapamycin effect *in vitro* on Schwann cell proliferation and neurotrophic factor production

Since Schwann cells (SC) were going to be used as supportive cells that would provide extracellular matrix and neurotrophic factors, it was necessary to demonstrate that rapamycin would not disrupt their function. Schwann cells (SCs) were isolated from the sciatic nerves of postnatal day four Fischer rat pups as previously described (Olson *et al*. 2009; Rooney *et al*. 2011) and placed in SC media (DMEM/F12 medium containing 10% fetal bovine serum, 100 units/mL penicillin/streptomycin (Gibco, Grand Island, New York, USA), 2 μM forskolin (Sigma, St. Louis, Missouri, USA), and 10 ng/mL recombinant human neuregulin-1-β1 extracellular domain (R&D Systems). After four passages, SCs were seeded at a density of 30,000 cells per well into 24 well plates with and treated with rapamycin doses ranging from 1 nM to 1 mM or ethanol vehicle in quadruplicate for 3 days at 37° C. SC proliferation was then measured using the CellTiter 96 AQueous One Solution Cell Proliferation MTS Assay kit (Promega) using the same protocol as for the fibroblast measurements.

To determine the concentration of brain-derived neurotrophic factor (BDNF) and glial-cell derived neurotrophic factor (GDNF) secreted from SCs with and without rapamycin treatment, 200 ul of supernatant cell culture media were also sampled after 3 days in culture for measurement by enzyme-linked immunosorbent assays (ELISA) (Human BDNF DuoSet and Human GDNF DuoSet ELISA kits (R&D Systems)). Growth factor concentrations were compared against a standard curve, according to the manufacturer’s protocol. Streptavidin-HRP conjugation was amplified for both BDNF and GDNF ELISAs using the ELAST biotinyl tyramide system (PerkinElmer, Waltham, Massachusetts, USA).

### Immunohistochemical characterization of cultured primary Schwann cells

The purity of primary SC cultures after four passages was determined by immunohistochemistry. 15,000 SCs were plated onto five 35 mm glass bottom culture dishes. Cells were washed with phosphate buffered saline (PBS), fixed with 4% paraformaldehyde, blocked with 7.5% bovine serum albumin (BSA), and incubated overnight at 4° C with rabbit anti-rat p75 neurotrophin receptor (p75(NTR) (Promega, 1:500) in PBS with 1% BSA. After washing in PBS with 0.1% triton X-100, cells were incubated with donkey anti-rabbit Cy3 conjugated secondary antibody (Millipore, 1:100) in PBS with 5% normal donkey serum and 0.3% triton X-100, and mounted in SlowFade Gold Antifade Reagent with 4',6-diamidino-2-phenylindole (DAPI) (Molecular Probes, Eugene, Oregon, USA) nuclear counterstain. Fifteen random fields (three from each plate) were imaged and analyzed using the Neurolucida software (MBF Bioscience, Williston, Vermont, USA). The number of cells positive for p75(NTR) staining was then counted as a percentage of the total number of cells counted with DAPI-stained nuclei.

### Poly-lactic-co-glycolic acid (PLGA)-rapamycin microsphere fabrication

A microsphere-based drug delivery strategy was used to provide sustained release of effective doses of rapamycin. Poly-lactic-co-glycolic acid (PLGA) microspheres were fabricated from polymer with a 50:50 lactic acid to glycolic acid ratio and 29 kDA molecular weight (5050 DLG 4A, Evonik Industries, Essen, Germany), using an oil-in-water emulsion-solvent evaporation technique, as previously described (Rooney *et al*. 2011; Rui *et al*. 2012). Microspheres containing different doses of rapamycin were constructed by dissolving 0.25 mg, 0.5 mg, 1 mg, or 4 mg of rapamycin in 100 μL of absolute ethanol. Each rapamycin solution was added dropwise and vortex emulsified in a solution of 250 mg PLGA in 1 ml of methylene chloride for 30 seconds. The mixture was re-emulsified for another 30 seconds in 2 ml 2% (w/v) poly(vinyl alcohol) (PVA), added to 100 ml of 0.3% (w/v) PVA solution and 100 ml of 2% (w/v) isopropyl alcohol, and stirred to evaporate methylene chloride for one hour. The microspheres were collected by centrifugation at 2000 rpm for 3 minutes and washed 3 times with distilled water. A microsphere aggregate was obtained for use in subsequent experiments by freezing at -80 °C for 1 h and vacuum drying overnight. Microspheres were fabricated under sterile conditions.

### Measurement of encapsulation efficiency of rapamycin within PLGA microspheres

The encapsulation efficiency of rapamycin within 0.25 mg, 0.5 mg and 1 mg per 250 mg PLGA loaded microspheres was determined by extracting the drug and quantitating its concentration against a standard curve. 5 mg of microspheres from each PLGA-rapamycin formulation were dissolved in 1 mL of acetonitrile. Samples were centrifuged at 3000 RPM to pellet remaining debris. 200 μL of each sample was transferred to a 96-well plate with UV transparent flat bottom and the absorbance at 278 nm was measured using a Spectra Max Plus 384 plate reader (Molecular Devices). These readings were compared to a standard curve prepared using known concentrations of free rapamycin dissolved in acetonitrile (0 μg/mL, 1 μg/mL, 2.5 μg/mL, 5 μg/mL, 10 μg/mL, 15 μg/mL, 20 μg/mL). The yield of rapamycin from each sample was compared to the amount that had been loaded initially into the microspheres to determine the percent encapsulation efficiency ((yield extracted/amount loaded) x 100) (Rui *et al*. 2012). All samples were processed in duplicate.

### Measurement of release kinetics of rapamycin from PLGA microspheres *in vitro*

10 mg of microspheres, from each rapamycin/PLGA dosage group were placed into 5 mL of PBS with 10% FBS and briefly vortexed. Five tubes were allocated to be incubated for 1, 2, 3, 4, or 5 weeks at 37° C for each formulation. Twice weekly a set of microspheres from each formulation group were pelleted (3500 RPM for 10 minutes), washed once with 5 mL of PBS and recentrifuged. After removal of the supernatant, 1 mL of acetonitrile was added to dissolve the microspheres through sonication and vortexing. Rapamycin yield from each sample was determined as in the loading protocol above and compared to the yield of corresponding fresh microspheres from the encapsulation efficiency experiments to determine the percentage of rapamycin remaining at each time point.

### Fabrication of OPF+ scaffolds containing rapamycin-releasing PLGA microspheres

Positively charged oligo[poly(ethylene glycol)fumarate] (OPF+) hydrogel scaffolds were fabricated as previously described (Chen *et al*. 2011; Dadsetan *et al*. 2009). The liquid polymer contained 1 g of OPF macromer dissolved in 0.05% (w/w) photoinitiator (Irgacure 2959, Ciba Specialty Chemicals, Tarrytown, New York, USA), 0.3 g N-vinyl pyrrolidinone (NVP) (Sigma) and 20% [2-(methacryloyloxy) ethyl]-trimethylammonium chloride (MAETAC) (Sigma) in 650 μL of deionized water. To make OPF+ scaffolds embedded with rapamycin-PLGA microspheres, 25 mg of microspheres of each rapamycin formulation were stirred into 250 μL of liquid OPF+ polymer solution. Approximately 35 μL of OPF+-PLGA microsphere solution was then cast by mold injection over seven parallel wires of 290 μm diameter spaced within a glass cylinder, and polymerized by exposure to UV light (365 nm) at an intensity of 8 mW/cm2 (Black-Ray Model 100AP). Individual scaffolds were cut into 2 mm lengths and sterilized prior to cell loading by immersion in serial dilutions of ethanol (40%, 70%, and 100% for 30 min each), followed by four changes of PBS. They were stored in PBS for 24 hrs at 4° C. The resulting scaffolds were 2.6 mm in diameter with internal channels measuring 450 μm in diameter. Each 2 mm multichannel OPF+ scaffold was calculated to be embedded with approximately 0.175 mg of microspheres.

### Analysis of fibroblast proliferation in vitro in response to rapamycin released from PLGA microspheres and OPF+ hydrogel scaffolds

30,000 NRK fibroblast cells per well were plated in 24 well tissue culture plates in DMEM with 10% FBS and incubated at 37° C in a 5% CO_2_ humidified atmosphere, then serum-starved and treated with TGF-β1 as described for the fibroblast proliferation assay above. 5 mg of rapamycin-containing PLGA microspheres (0.25 mg, 0.5 mg or 1 mg rapamycin per 250 mg PLGA), or two 2 mm multichannel scaffolds embedded with microspheres, were co-cultured with fibroblasts within Millicell hanging cell culture inserts (Millipore, Billerica, Massachusetts, USA). To allow for variable durations of drug release prior to the physiologic assay, PLGA microspheres were incubated in hanging cell culture inserts at 37° C in DMEM with 10% FBS culture media, which was changed daily for 1, 2, 3, or 4 weeks, while OPF+ scaffolds embedded with microspheres were incubated with daily media changes for 1 week. The inserts were then transferred to four wells containing 30,000 fibroblasts per well as described above and co-cultured for 2 days. The MTS assay to measure TGF-β1 stimulated fibroblast proliferation was then carried out as described above.

### Scaffold loading

OPF+ scaffolds incorporating PLGA microspheres were loaded with 8 μL of Matrigel containing 10^5^ cells/μL Schwann cells (SC) or Matrigel (MG) alone as previously described (Chen *et al*. 2011; Hakim *et al*. 2015). The loaded scaffolds were then incubated in Schwann cell media for 24 hours at 37° C in 5% CO2 before implantation into animals.

### Animal allocation for spinal cord implantation of OPF+ scaffolds embedded with rapamycin-PLGA microspheres

A total of 83 adult female Fischer rats (Harlan Laboratories, Indianapolis, Indiana, USA) weighing approximately 200 g were used in this study, with 53 animals used in the first cohort of animals and 30 in the replication cohort. 75 rats survived the entire length of the experiment, with 6 early animal deaths in the first cohort and 2 early animal deaths in the second cohort due to complications during surgeries or within the week after surgery. In the first cohort, animals were designated to receive OPF+ scaffold implantation with PLGA microspheres following complete spinal cord transection at the T9 level. Scaffold conditions varied as to the inclusion of primary Schwann cells, and as to the dosage of rapamycin release (Table 1). There was a 6 week endpoint in the first experimental cohort. Weekly locomotor scores were assessed and following which scaffolds were embedded and sectioned in the transverse or longitudinal plane at their midpoint for Masson’s trichrome staining and immunohistochemical studies.

**Table 1.**
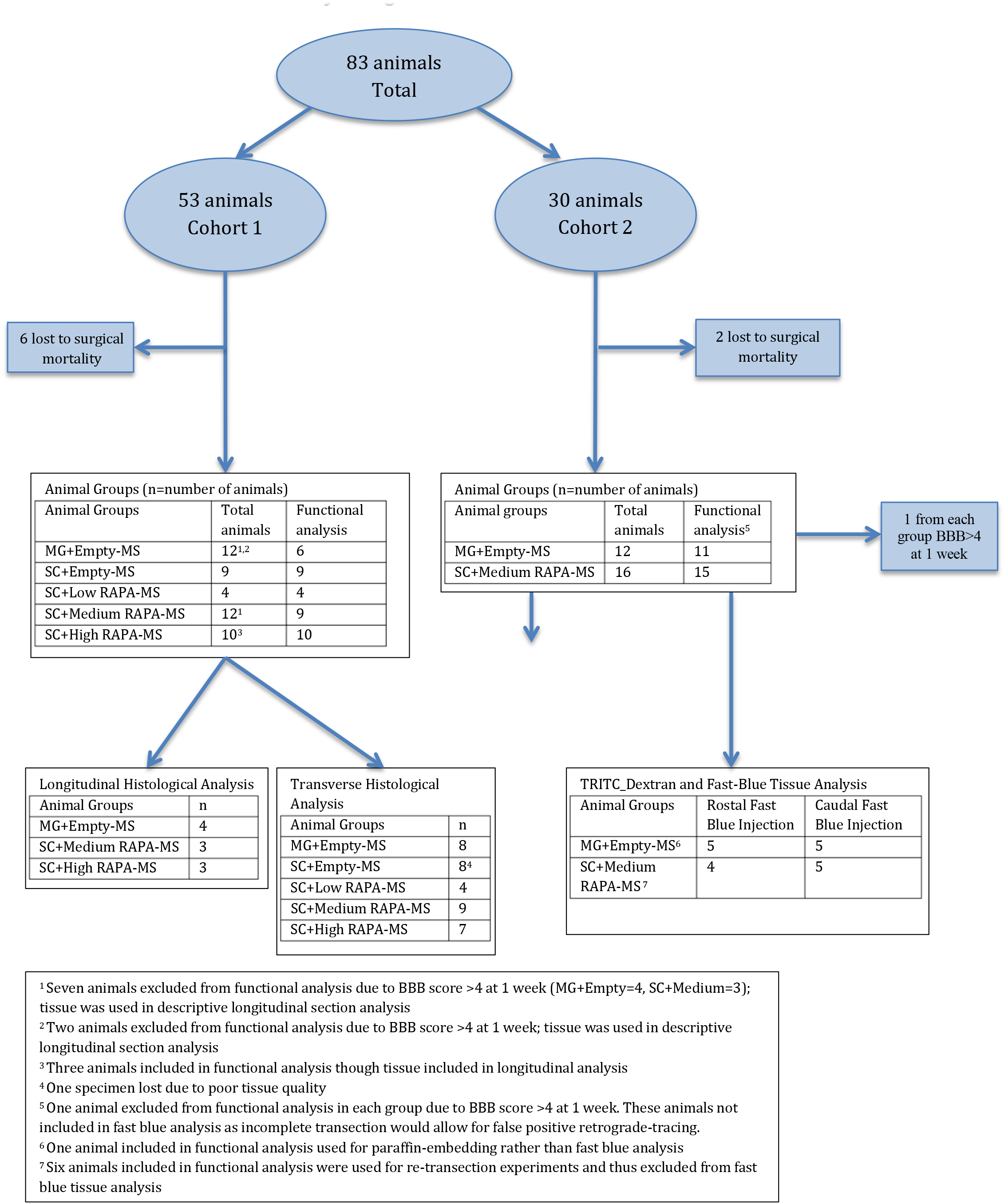
Overview of Animal Study Design

Thirty additional animals were used in a second cohort with a 6 week endpoint (Table 1). Two animals died during surgery, leaving 28 animals for the cohort. This was designed to replicate the functional outcome findings following transplantation of OFP+ scaffolds with Schwann cell and medium dose rapamycin microspheres (1 mg per 250 mg PLGA) compared to Matrigel only scaffolds with empty microspheres. These animals were also designated for retrograde fast blue injection studies, spinal cord retransection, somatosensory evoked potential neurophysiology, and vascular studies.

### Animal care, spinal cord transection and scaffold implantation surgeries

All animal procedures were performed according to the guidelines of the Mayo Clinic Institutional Animal Care and Use Committee (IACUC). Animals were kept on a standard 12 hour light-dark cycle with access to food and water *ad libitum* in conventional housing in accordance with National Institutes of Health (NIH) and U.S. Department of Agriculture guidelines. Animal spinal cord transection surgical techniques, scaffold implantation and post-operative care were as we have previously described (Hakim *et al*. 2015). Animals received perioperative analgesia for 1 week using oral acetaminophen (Mapap, Major Pharmaceutiacals, Livonia, Michigan, USA), oral Baytril (Bayer Corporation, Shawnee, Kansas, USA) and subcutaneous Buprenex (Reckitt Benckiser Pharmaceuticals Inc, Richmond, Virginia, USA). Animals were anesthetized for surgery with intraperitoneal ketamine (Fort Dodge Animal Health, Fort Dodge, Iowa, USA) and xylazine (Lloyd Laboratories, Shenandoah, Iowa, USA). Animals were randomly assigned to experimental groups and surgeries were performed by a surgeon blinded to animal groups using our previously described approach (Hakim *et al*. 2015; Madigan *et al*. 2014). Laminectomy was performed at the T9-T10 vertebral level and the spinal cord was completely transected using a number 11 scalpel blade. Complete transection was verified with a hooked probe. After transection, the two cut ends of the spinal cord retracted to form a two mm gap. Prepared OPF+ scaffolds were implanted between the free ends of the spinal cord, with the scaffold channels oriented parallel to the length of the spinal cord. After surgery animals were kept in low-walled cages to allow easy access to food and water. Bladders were expressed three times daily after surgery and animals were given antibiotics and analgesics as necessary. All rats were cared for 24/7 by veterinarians and technicians with experience in the management of rat spinal cord injury.

### Open field hind limb locomotor function testing

Hind limb function of scaffold-implanted animals was evaluated weekly through open field testing by two to four independent observers blinded to the animal groups and scored using the 21 point Basso, Beattie, and Bresnahan (BBB) locomotor rating scale (Basso *et al*. 1995). The BBB score for each left and right hind limb was obtained by averaging the scores given by the independent observers. Scores for the left and right hind limbs were then averaged to obtain the overall BBB score for each animal, which was used for statistical analyses. As in previous our previous studies (Chen *et al*. 2017; Hakim *et al*. 2015) any animal with a BBB score greater than 4 (representing slight movement of all three joints of the hind limb) at one week post implantation was considered to have a physiologically incomplete transection and was excluded from further functional analysis. A total of 10 rats distributed across all groups from both cohorts were excluded in this way (Table 1). Spinal cords of the excluded animals in the first cohort were processed and included in subsequent histological analysis.

### Tibial SSEP monitoring

Somatosensory evoked potentials (SSEPs) were recorded from animals in the second cohort through subcutaneous scalp needle electrodes (Technomed Europe, Maastricht, Netherlands) following bilateral tibial nerve stimulation using a Nicolet Viking IV system (Viasys Healthcare, Conshohocken, Pennsylvania, USA) as previously described (Cloud *et al*. 2012). Baseline SSEPs were recorded bilaterally before scaffold implantation surgery, and post implantation at 1, 6, and 7 weeks (in 6 retransection animals). The SSEP waveforms were analyzed by a clinical neurophysiologist who was blinded to animal group and the time point of recording. Numerical scores were given to the waveforms according to the following descriptions: absent (1), likely absent (2), likely present (3), and present (4). The scores for waveforms obtained through stimulation of the left and right hind limbs were averaged for each animal to obtain an overall SSEP score, which was then used for statistical analysis.

### Fast Blue retrograde tracer dye injection and spinal cord retransection surgeries

Ten animals from both groups of the second cohort underwent a second surgery 5 weeks after scaffold implantation for Fast Blue (FB) retrograde tracer analysis, as we have previously described (Chen *et al*. 2009; Chen *et al*. 2017). The spinal cord was re-exposed at the level of the prior implantation and 0.6 μL of Fast Blue dye was stereotactically injected into the spinal cord with a Hamilton syringe through a second surgical laminectomy at a distance of 5 mm rostral (5 animals/group) or caudal (5 animals/group) to the implanted scaffold. Fast Blue dye was slowly injected over a period of 30 seconds, and the needle was left in place for another 30 seconds to prevent dye leakage. The muscle and skin was then sutured closed as described for the transection surgery. Animals were allowed to recover for an additional 7 days before tissue harvest and analysis. Since it was important to demonstrate that functional recovery was due to axonal regeneration, six animals in the medium dose group of the second cohort underwent a second surgery for spinal cord retransection at week 6. The spinal cord was re-exposed at the original laminectomy site and the implanted scaffold was visualized. The laminectomy site was slightly expanded rostrally to allow for a retransection of the spinal cord immediately adjacent to the scaffold using a No. 11 scalpel. Complete transection was verified with a hooked probe. The muscle and skin was then sutured closed as described for the transection surgery.

### TRITC-dextran tail vein injections

TRITC-dextran infusion was used to analyze vascular perfusion in the regenerating tissue. A tail vein injection of 250 μL of a 0.9% saline solution containing 10 mg/mL TRITC-dextran (average molecular weight 65000-85000 Da, Sigma) was performed on the animals in the second cohort with Fast Blue dye injections two hours before sacrifice at week 6. The animals were warmed under a heat lamp until the tail vein dilated and were placed in a plastic restraint device. The tail was scrubbed with ethanol, one vein was selected and 250 μL of fluid was injected via a 27 gauge needle. The needle was then removed and pressure was applied to the tail using sterile gauze for 30 seconds. Tissue analysis of TRITC-dextran is described below.

### Tissue preparation and sectioning

At the conclusion of the final functional studies, animals were humanely euthanized with an intraperitoneal injection of 0.4 mL sodium pentobarbital (40 mg/kg) (Fort Dodge Animal Health, Fort Dodge, Iowa, USA). For paraffin tissue embedding and sectioning, spinal cord segments were prepared from animals following transcardial perfusion with 4% paraformaldehyde in PBS as previously described (Hakim *et al*. 2015). For frozen sectioning of Fast Blue dye injected animals with tail vein injections of TRITC-dextran, the vertebral column with spinal cord was removed *en bloc* without animal perfusion and post fixed for 4 days in 4% paraformaldehyde with 10% sucrose at 4° C. The spinal cord was then dissected from the vertebral column and cryoprotected in 30% sucrose at 4° C for 24 hours before being processed for cryostat embedding. The spinal cord length was cut sectioned into 1.5 cm segments designated P1, P2 and P3 moving rostrally, and S1 and S2 moving caudally. The P1 segment contained the OPF+ scaffold. These segments were then embedded with Tissue Freezing Medium (TFM) (Triangle Biomedical Sciences, Durham, North Carolina, USA), cut longitudinally into 30 μm sections (Reichart HistoSTS Cryostat Microtome) and mounted on numbered slides with Aqua-Mount (ThermoFisher).

### Analysis of collagen scarring within channels

Slides with transverse tissue sections were selected from the central portion of the scaffolds for staining with the Masson trichrome kit (ThermoFisher). Acquired images were analyzed using the Neurolucida software (MBF Bioscience, Williston, Vermont, USA). A digital grid with 100 μm divisions was overlaid on the tissue, and digital markers for collagen scar or unscarred tissue were placed at each grid intersection within the scaffold channels. The area of collagen scar and unscarred tissue within scaffold channels was then calculated using the Cavalieri probe in the Stereo Investigator software (MBF Bioscience), as previously described (Hakim *et al*. 2015)

### Immunohistochemistry

Paraffin transverse sections were selected for immunohistochemistry from the central portion of the scaffolds, adjacent to the Masson trichrome stained slides. Sections were deparaffinized with xylene, rehydrated with graded ethanol and rinsed with distilled water. Heat mediated antigen retrieval was performed by incubating the sections in 1 mM EDTA in PBS for 30 min in a rice steamer. After blocking with 10% normal donkey serum (NDS) in PBS for 30 min sections were incubated overnight at 4° C in primary antibody diluted in 5% NDS with 0.3% Triton X-100 in PBS. Sections were washed with 0.1% Triton X-100 in PBS before incubating for one hour in secondary antibody diluted in 5% NDS with 0.3% Triton X-100 in PBS. After washing with 0.1% Triton X-100 in PBS, sections were mounted with SlowFade Gold Antifade Reagent with DAPI (Molecular Probes, Eugene, Oregon, USA). For DAB (3,3’-diaminobenzidine) staining with nickel enhancement, the DAB substrate kit (Vector Laboratories, Burlingame, California, USA) was used according to the manufacturer’s instructions and slides mounted with Aqua-Mount.

Primary antibodies were used against platelet derived growth factor receptor β (PDGFRβ) (Rabbit anti-rat, 1:50, Cell Signaling Technology, Danvers, Massachusetts, USA); rat endothelial cell antigen-1 (RECA-1) (Mouse anti-rat, 1:100, Abcam, Cambridge, Massachussets, USA); p75(NTR) (Rabbit anti-rat, 1:800 Promega) and collagen IV (rabbit anti-rat, 1:300 (Abcam)). Secondary antibodies included Alexafluor 647-conjugated donkey Anti-Rabbit IgG (1:200, Jackson ImmunoResearch, West Grove, Pennsylvania, USA); Cy3-conjugated donkey anti-rabbit IgG antibody (1:200, Millipore, Billerica, Massachusetts, USA); Cy3-conjugated donkey anti-mouse IgG (1:200, Millipore), and horseradish peroxidase (HRP)-conjugated donkey anti-rabbit IgG (1:200, Millipore).

### Analysis of Schwann cell number and phenotype *in vivo*

For Schwann cell *in vivo* analysis, transverse sections were selected from the central portion of the scaffolds for staining with p75(NTR) to identify Schwann cells. Automatic object detection was then used in the Neurolucida software (MBF Bioscience, Williston, Vermont, USA) to draw contours around the p75(NTR) stained Schwann cells within the scaffold channels. The sum of the contoured areas outlined within the scaffold channels with automatic object detection was then used for analysis as previously described (Hakim *et al*. 2015).

### Microscopy

Images used for Masson Trichrome and immunohistochemical analysis were acquired on a modified Zeiss Axioimager A-1 microscope (Zeiss, Thornwood, New York, USA) equipped with a motorized specimen stage for automated sampling (Ludl Electronics; Hawthorne, New York, USA) and a QICAM 12-bit Color Fast 1394 camera (QImaging, Surrey, British Columbia, Canada). FB-labelled neurons were images using C-Apochromat 10x and 20 x objective lenses on a Zeiss LSM510 laser scanning confocal microscope. Additional confocal images were acquired using a LSM 780 confocal microscope (Zeiss, Thornwood, New York, USA).

### Sterological analysis of collagen IV stained blood vessels

Surface density (Sv) and length density (Lv) of bloods vessels were calculated using unbiased stereology by applying linear and plane grids, respectively, to count vessels as previously described (Madigan *et al*. 2014). Mean vessel diameter (d) was calculated from the ratio of surface to length density, according to the equation d = Sv/(Lv x π).

### Quantification of endothelial and perivascular cell coverage of vascular structures

Fiji image analysis software (Schindelin *et al*. 2012) was used to identify the endothelial and perivascular cell coverage of each individual vessel within scaffold channels. A high definition interactive pen display, the Cintiq 24HD touch (Wacom Technology Corporation, Vancouver, Washington, USA) was used by an observer blinded to animal group to draw a contour along the pericyte and endothelial cell staining surrounding the lumen of the vessel. Each contour was labeled and measured in length. After calculating a pericyte to endothelial cell ratio for every individual vessel within a channel, the ratios were divided into 5 clusters according to the extent of pericyte coverage. i.e. <0.75, 0.75-1.5, 1.5-2.25, 2.25-3.0, 3.0-3.75, 3.75-7.0, or >7.0. After quantifying how many vessels of each cluster were present in each channel, a mean number of vessels per channel were calculated for each animal. Animal groups were compared using a two-way analysis of variance (ANOVA). Posthoc analysis was performed by Newman-Keuls multiple comparison method.

### Analysis of vascular function using TRITC-dextran

In the animals with TRITC-dextran tail vein injection, quantification of the amount of intravascular TRITC dye was performed by measuring the surface areas of red fluorescence within longitudinal image fields. Images were systematically taken at 20x magnification spanning the width of the central scaffold channel only of the seven-channel scaffolds to ensure consistency of sampling. The total surface area of intravascular TRITC dye was calculated by setting a defined sensitivity for object detection in the stereology software, which was kept constant across all sections analyzed. Automatic object detection was used for this quantification in the Neurolucida software suite (MBF Bioscience) by manually drawing contours around intense TRITC dye foci within using the same settings of 70.2% sensitivity and highest point density for all animals. The output from object detection software listed the individual surface areas for each fluorescent object, which were summed for each animal and then analyzed as mean area per group.

### Quantification of Fast Blue dye labeled regenerated neurons

Fast Blue (FB)-labeled neurons were quantified as previously described (Chen *et al*. 2009; Chen *et al*. 2017) within 30 micron thick longitudinal frozen sections. Labeled neurons in the P1-P3 and S1-S2 (Fig. 6 A, B) segments that were clearly identified as having a visible nucleus and typical cell body morphology were counted within every other section by observers blinded to the animal groups. 158 nuclear diameters were measured in their longest dimension in FB-labeled neurons using NIH ImageJ software in random spinal cord segment images from both animal groups. The mean nuclear diameter (7.43 +/-0.16 microns) was then used in Abercrombie’s formula (Abercrombie 1946; Guillery 2002) for FB-neuron count correction, to account for the number of neuronal profiles divided into two by the microtome and thus present in two adjacent sections. The formula provides a ratio of the estimated number of nuclei to the observed number, T/T+h, where ‘T’ is the section thickness and ‘h’ is the mean nuclear diameter. A calculated correction factor of 0.802 was then derived and applied to the observed counted profile number in each spinal cord segment, indicating that the overestimation of counts was 24.7%. Neuron counts from each 1.5 cm spinal cord segment in the rostral cord (P1+P2+P3) after caudal dye injection and in the caudal cord (S1+S2) after rostral dye injection were then summated to obtain the total number of regenerating neurons. Neurons counted within the P1 segment, which was longitudinally bisected by the OPF+ scaffold, were separated into rostral and caudal counts.

### Statistical analysis

All analyses were performed by observers blinded to treatment groups. The averages for proliferation of fibroblasts and Schwann cells, Schwann cell BDNF and GDNF production, DRG neurite outgrowth, percent channel area without collagen scarring, channel blood vessel length, channel blood vessel surface area, channel blood vessel mean diameter, and hind limb function in the subset of animals that received retransections were all analyzed by one-way ANOVA with *post hoc* analysis using Newman-Keuls multiple comparison test. The average BBB scores for hind limb function in the first and second cohorts and pericyte to endothelial cell ratios of scaffold blood vessels were analyzed by two-way repeated measures ANOVA with *post hoc* analysis using Newman-Keuls multiple comparison test. Rostral and caudal total Fast Blue labeled neuron counts and areas of TRITC dye foci were analyzed using the nonparametric Mann-Whitney test. Statistical analyses were performed using GraphPad Prism 6 software (GraphPad Software Inc., San Diego, California, USA). All data are presented as mean ± standard error of the mean (SEM). Values of p < 0.05 were considered statistically significant.

## Results

### Rapamycin inhibits fibroblast and Schwann cell proliferation and stimulates neurotrophin production

The anti-fibrotic dose range for rapamycin was established using a proliferation assay with rat kidney (NRK) fibroblasts treated with 5 ng/ml recombinant human transforming growth factor (TGF)-β1 (Supplementary Fig. 1 A). Rapamycin inhibited fibroblast proliferation between 1 nM and 100 nM without affecting cell viability. Neurite outgrowth from dorsal root ganglion explants in 1 nM to 100 nM rapamycin was unaffected by drug treatment after 24 and 48 hours (Supplementary Fig. 1 B and C). Doses of 1 nM to 1 mM rapamycin inhibited proliferation of primary rat SC *in vitro* over 3 days (Supplementary Fig. 1 D). BDNF concentration was increased at 80 nM rapamycin and GDNF at 1 mM rapamycin (Supplementary Fig. 1 E and F). Primary SC were used after four passages to simulate cells loaded into hydrogel scaffolds for *in vivo* studies. 90.4 +/-0.5% of these SCs maintained p75(NTR) expression (Supplementary Fig. 2).

### Rapamycin released from PLGA microspheres and OPF+ scaffolds inhibits fibroblast proliferation

Poly-lactic-co-glycolic acid (PLGA) microspheres were fabricated to target sustained release of rapamycin at doses established above. PLGA microspheres contained 0.25 mg, 0.5 mg, 1 mg or 4 mg of rapamycin per 250 mg PLGA. Encapsulation efficiency was 104 ± 9%, 100 ± 10%, and 107 ± 8%, respectively. Kinetics of drug release were determined over 5 weeks. (Supplementary Fig. 3 A).

Five micrograms of the 0.25 mg, 0.5 mg and 1 mg rapamycin per 250 mg PLGA microspheres inhibited NRK fibroblast proliferation over 4 weeks (Supplemental Fig. 3 B). TGF-β1-stimulated fibroblast proliferation was prevented *in vitro* when the same three formulations of rapamycin-PLGA microspheres were incorporated into the oligo[poly(ethylene glycol) fumarate] (OPF+) hydrogel scaffolds at a density of 1 mg microspheres per 10 μl liquid OPF+ in 2 mm length multichannel scaffolds (Supplementary Fig. 3 C). OPF+ scaffolds embedded with 0.25 mg, 1 mg and 4 mg rapamycin per 250 mg PLGA were used in the animal studies.

### Rapamycin from PLGA microspheres in OPF+ scaffolds reduces fibrosis after SCI

OPF+ scaffolds were implanted into five groups of Fischer rats after spinal cord transection at the T9 level (Table 1). The first cohort of 53 animals was used for a rapamycin dose response study with scaffolds containing primary SC compared to SC-loaded scaffolds with empty microspheres and Matrigel scaffolds with empty microspheres. Collagen deposition within scaffold channels after 6 weeks and weekly locomotor scores were measured. All animals were sacrificed at 6 weeks post implantation for tissue sectioning and analysis (Table 1).

At 6 weeks scaffolds containing empty PLGA microspheres, with or without SC developed dense collagen scarring throughout channels (Fig. 1 A and B). Low dose rapamycin animals were similar (Fig. 1 C). Both medium and high dose rapamycin had significantly less channel collagen deposition (Fig. 1 D and E). 11.54 ± 0.91 % and 10.77 ± 1.38 % of the core channel area remained clear of collagen in the Matrigel-only (n=8) and SC-loaded scaffolds with empty microspheres (n=8) groups, with 8.01 ± 2.40%, 50.35 ± 8.45%, and 62.04 ±2.85% of the channel area clear in the SC-loaded scaffolds with low dose (n=4), medium dose (n=9) and high dose rapamycin (n=7) groups, respectively (Fig. 1 F).

**Fig 1.**
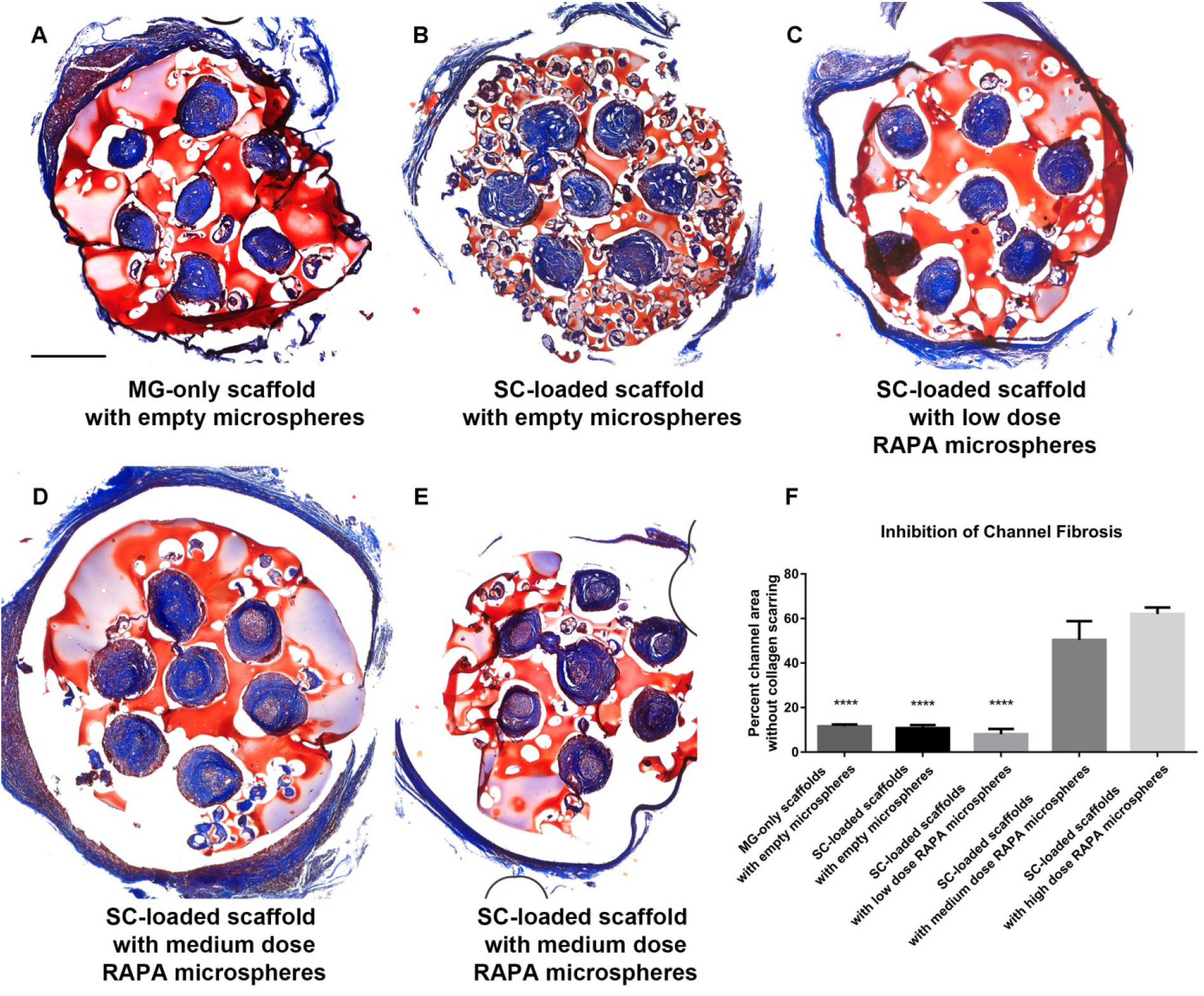
Inhibition of collagen scarring within scaffold channels by rapamycin *in vivo*. (A) Masson trichrome staining from the midpoint of OPF+ scaffolds demonstrated high levels of collagen scarring (blue) in the channels of animals implanted with MG-only scaffolds with empty microspheres. (B) SC-loaded scaffolds with empty microspheres. (C) SC-loaded scaffolds with low dose rapamycin microspheres. (D) In contrast, a significant amount of unscarred tissue (pink) was observed within the channel centers of animals implanted with SC-loaded scaffolds with medium dose rapamycin microspheres (E) SC-loaded scaffolds with high dose rapamycin microspheres. (F) Percent channel area without collagen scarring in each of these groups was quantitated. (****p<0.0001) denotes significant decrease from both the SC-loaded scaffolds with medium dose rapamycin microspheres and SC-loaded scaffolds with high dose rapamycin microspheres groups. 500 urn scale bar in panel (A) applies to panels (B) through (E).

### Rapamycin influences Schwann cell number and phenotype *in vivo*

SCs were identified by p75(NTR) staining in transverse sections through the central portion of the scaffolds (Supplementary Fig. 4 A-E). There was an increased area of p75(NTR) staining in animals implanted with SC-loaded scaffolds with empty microspheres compared to other groups (Supplementary Fig. 4 F). The trend towards reduction in SC numbers at 6 weeks post implantation in the rapamycin-treated groups is in agreement with the *in vitro* data showing inhibition of SC proliferation by rapamycin. SCs in rapamycin-treated channels exhibited a more differentiated, spindle shaped morphology compared to the large amoeboid morphology seen in channels without rapamycin.

### Rapamycin promotes functional recovery in two separate animal cohorts

We next determined the effect of rapamycin treatment on functional locomotor recovery in the first cohort (Table 1). After 4 weeks, rats implanted with SC-loaded scaffolds containing medium dose rapamycin microspheres (SC+Medium RAPA-MS) demonstrated improved functional recovery (BBB score 4.69 ± 0.57, n=9 animals) when compared to animals implanted with Matrigel-only scaffolds with empty microspheres (1.58 ± 0.57, n=6) (p=0.0059) (MG+Empty-MS; Fig. 2 A). After 5 weeks, the mean score in the medium dose rapamycin group was 5.89 ± 0.45, further improved over the MG+Empty-MS group (1.83 ± 0.52)(p=0.0001) and the SC-loaded group with empty microspheres (SC+Empty-MS; 2.75 ± 0.75)(p=0.0012). At week 6, the mean score in SC+Medium RAPA-MS was 5.72 ± 0.53, compared with MG+Empty-MS (1.17 ± 0.25; p<0.0001), and SC+Empty-MS (3.08 ± 0.0.86; p=0.0016). To replicate these results a second cohort of 28 animals (Table 1) was used; SC+Medium RAPA-MS (1 mg rapamycin per 250 mg PLGA; n=16 animals), and MG+Empty-MS (n=12 animals). Blinded weekly measurement of BBB locomotor function demonstrated divergence of functional recovery at the 4-week time point (Fig. 2 B). The mean BBB score in the SC+Medium RAPA-MS group (3.16 ± 0.52) (n=15) was higher than that measured in the MG+Empty-MS (1.06 ± 0.25)(n=11)(p<0.01). At week 6 the mean score in the SC+ Medium RAPA-MS group was 4.91 ± 0.46 compared to the MG+Empty-MS group (3.23 ± 0.58) (p<0.05).

**Fig. 2.**
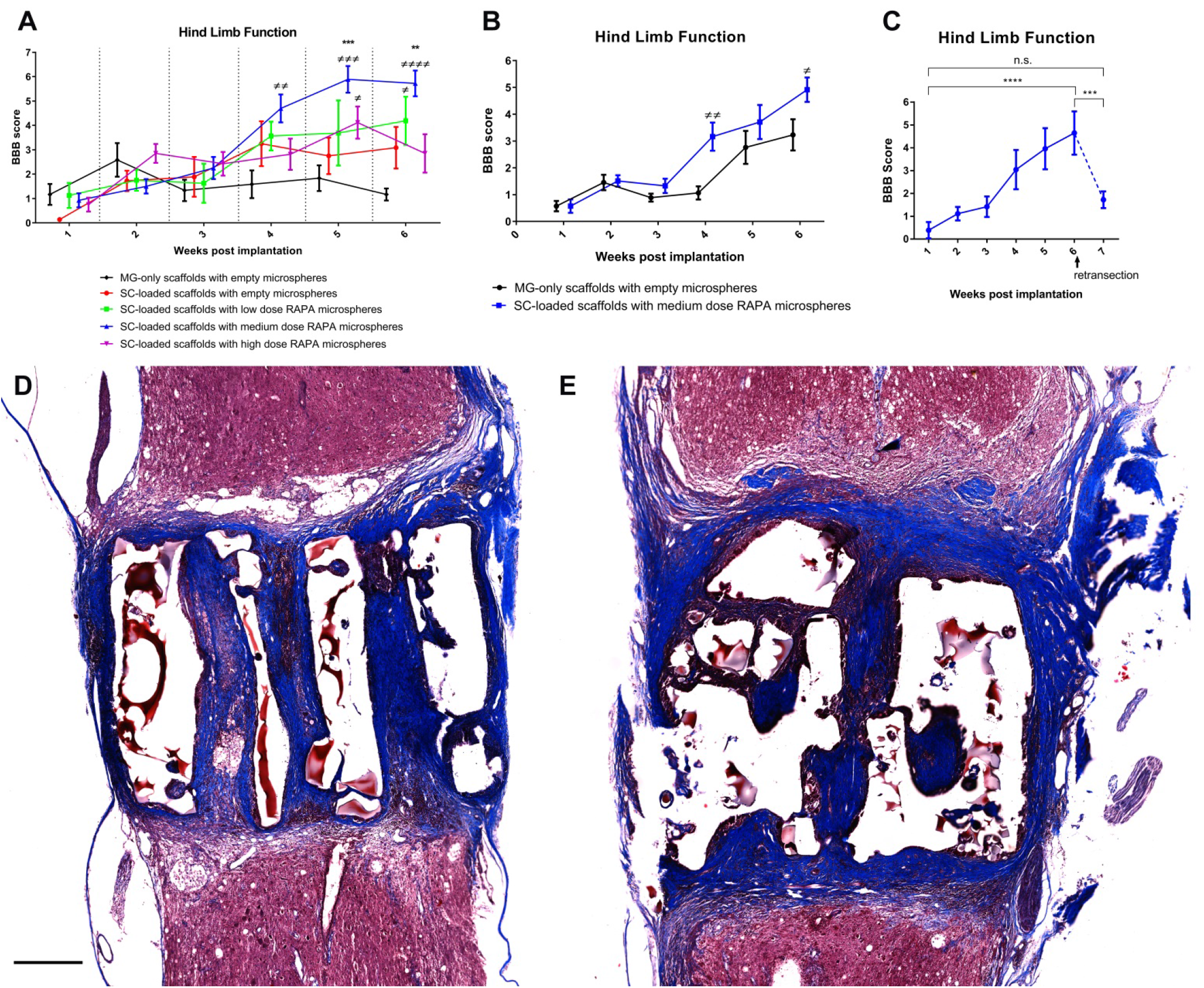
BBB scores in two animal cohorts, including spinal cord retransection. (A) Hind limb recovery over time in open field testing using the BBB scale in cohort 1. (B) Second cohort of scaffold-implanted animals and with SC and medium dose rapamycin. (C) Scaffold implanted animals with retransection at 6 weeks post implantation. Panel A (≠) denotes significant difference from MG-only scaffolds with empty microspheres group while (*) denotes significant difference from both MG-only scaffolds with empty microspheres (p=0.0001 week 5, p<0.0001 week 6) and SC-loaded scaffolds with empty microspheres groups (p=0.0012 week 5 and p=0.0016 week 6). At week 6, a significant difference was measured between the SC scaffolds with medium and high rapamycin doses (p=0.029). In panel B, (*) denotes significant difference from MG-only scaffolds with empty microspheres group (p<0.01 at week 4 and p<0.05 at week 6). (D) Longitudinal section through the center of implanted SC-loaded OPF+ scaffolds with medium dose rapamycin microspheres stained with Masson trichrome at 6 weeks post implantation. (E) Similar longitudinal section from an animal 7 weeks post implantation following retransection one week earlier. Longitudinal section (rostral end upward) confirms disruption of the scaffold tissue and channel architecture. Scale bar 500 μm (A and B).

To confirm that functional improvement was related to regeneration through the implanted scaffold, 6 animals from the SC+Medium RAPA-MS group underwent spinal cord retransection at the rostral tissue-scaffold interface at 6 weeks post implantation. One week later BBB scores were 1.72 ± 0.36 compared to 4.64 ± 0.95 (p = 0.011) at week 6 (Fig. 2 C). Histological assessment of the retransection site confirmed that compared to week 6 animals without retransection (Fig. 2 D), week 7 animals showed tissue disruption at the rostral tissue-scaffold interface with a characteristic fibrinoid scar (arrow) resembling that observed at week 1 post transection injury as previously reported (Hakim *et al*. 2015) (Fig. 2 E).

### Somatosensory evoked potential measurement

Pre-operative waveforms were present in all animals at baseline. SSEP waveforms were scored to be absent or likely absent (mean scores of 2.045 ± 0.142 in MG+Empty-MS and 1.87 ± 0.11 in SC+Medium RAPA-MS) at 1 week post implantation, confirming the complete nature of the transection injury. Definitive restoration of SSEP waveforms was not observed in either group at 6 weeks post implantation, and no change in the scoring was seen between week 6 and week 7 retransection measurements. (Supplementary Fig. 5).

### Rapamycin and Schwann cells influence vascularization, vessel architecture and function

Vascularity of scaffold channels was assessed by collagen IV staining of blood vessel basement membranes in transverse sections through the central portion of the scaffolds in the first cohort of animals (Fig. 3 A-E). Blood vessel length (a sum function of the number of blood vessels), mean vessel surface area, and mean diameter were analyzed through unbiased stereology (Fig. 3 F-H). SC+Empty-MS had a higher vessel length (482.5 ± 57.09 μm) and vessel surface area (12,034 ±775.6 μm^2^) than MG+Empty-MS (192.2 ± 27.61 μm and 3,341 ± 516.3 μm^2^, respectively) (p<0.05 for each comparison). Blood vessel length in the SC+Low RAPA-MS (626.6 ± 104.9 μm) and SC+Medium RAPA-MS (475.3 ± 103.8 μm) groups were statistically equivalent to the SC+Empty-MS group. Vessel surface area was similarly equivalent in the SC+Low RAPA-MS (12,983 ±2,397 μm^2^) and SC+Medium RAPA-MS (9399 ± 2203 μm^2^) groups compared to the SC+Empty-MS group. The presence of Schwann cells with no rapamycin or with low and medium rapamycin positively influenced blood vessel formation in OPF+ channels. However, vessel length (168.0 ± 53.96 μm^2^) and surface area (3,762 ± 1,379 μm^2^) in SC-loaded scaffolds eluting the high dose of rapamycin was equivalent to Matrigel-only scaffolds without rapamycin.

**Fig. 3.**
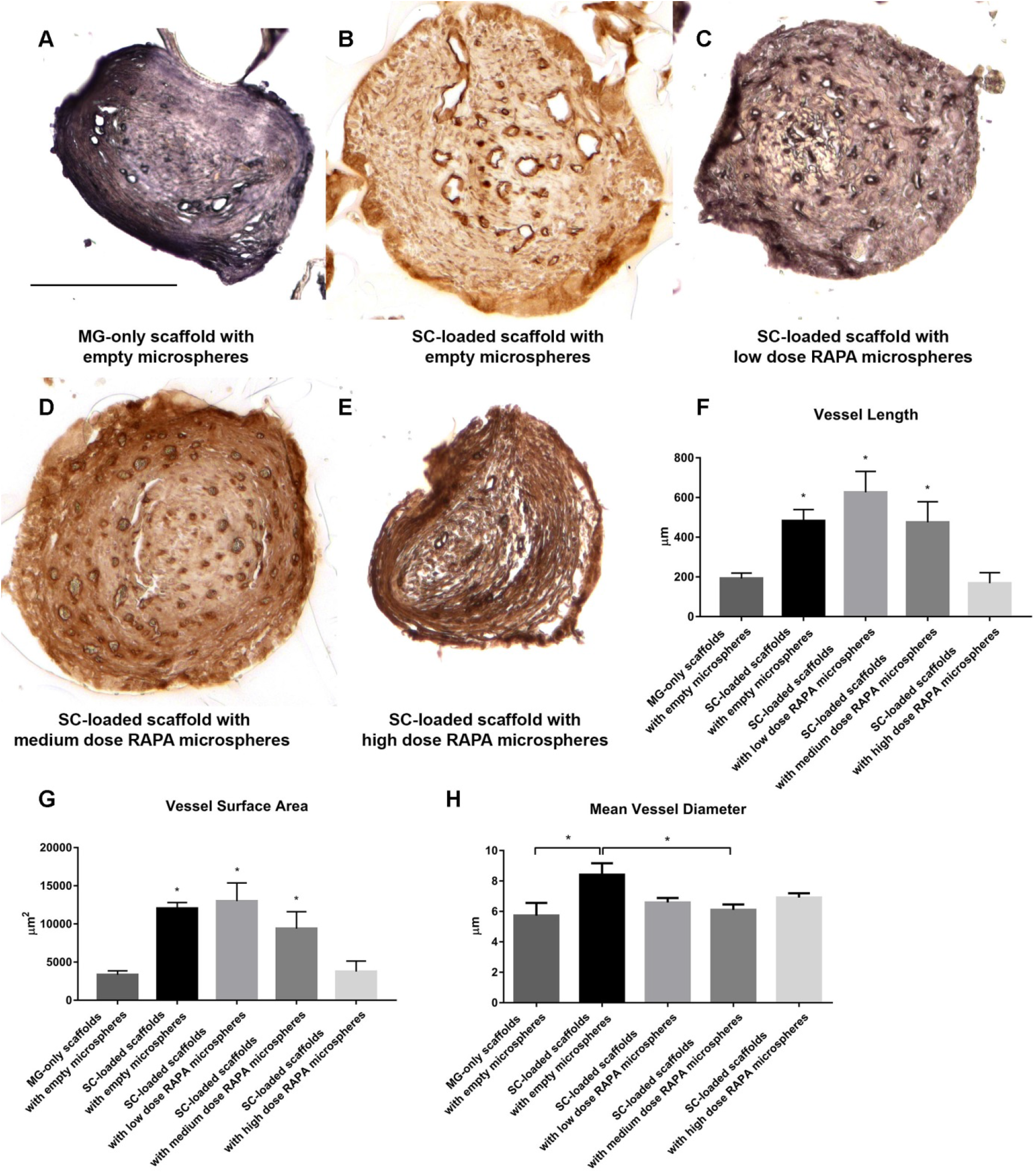
Vascularity and vessel morphology within scaffold channels in vivo. Images of individual scaffold channels stained for blood vessel basement membrane with collagen IV. (A) Few vessels in MG-only scaffolds with empty microspheres. (B) Large irregular vessels in SC-loaded scaffolds with empty microspheres. (C, D) Many smaller diameter vessels in SC-loaded scaffolds with low dose and medium dose rapamycin microspheres. (E) Few vessels in SC-loaded scaffolds with high dose rapamycin microspheres. 200 μm scale bar (A-E). (F) Quantitative stereological measurements of vessel length. (G) Stereological measurement of vessel surface area. (H) Quantitative stereological measurements of mean vessel diameter. In panels (F) and (G), (*) denotes significant difference from both MG-only scaffolds with empty microspheres and SC-loaded scaffolds with high dose rapamycin microspheres groups (p<0.05). In panel (H), *p<0.05 is denoted.

Blood vessel morphology also differed in different conditions. Vessels within SC+Empty-MS scaffolds had an irregular shape and larger mean diameter in contrast to smaller diameter vessels observed in animals with Matrigel-only scaffolds and the rapamycin treated groups (Fig. 3 A-E). The mean blood vessel diameter in SC+Empty-MS was 8.413 ± 0.76 μm, in Matrigel-only scaffolds 5.728 ± 0.84 μm (p<0.05), and in SC+Medium RAPA-MS was 6.10 ± 0.36 μm (p<0.05) (Fig. 3 H). Blood vessel diameters in the low and high dose rapamycin groups were also smaller than SC-loaded scaffolds with empty microspheres. Cellular structure of vessel walls varied between conditions. Small diameter vessels with normal endothelial cell staining (RECA-1) and pericyte (PDGFRβ) coverage were consistently observed in the channels of animals implanted with SC+Medium RAPA-MS scaffolds (Fig. 4 A). Irregularly bordered vessels with hypertrophic pericytes (Fig. 4 B) and vessels composed only of pericytes without endothelial cells (Fig. 4 C) were observed in SC+Empty-MS animals consistent with incomplete or irregular vessel formation.

**Fig. 4.**
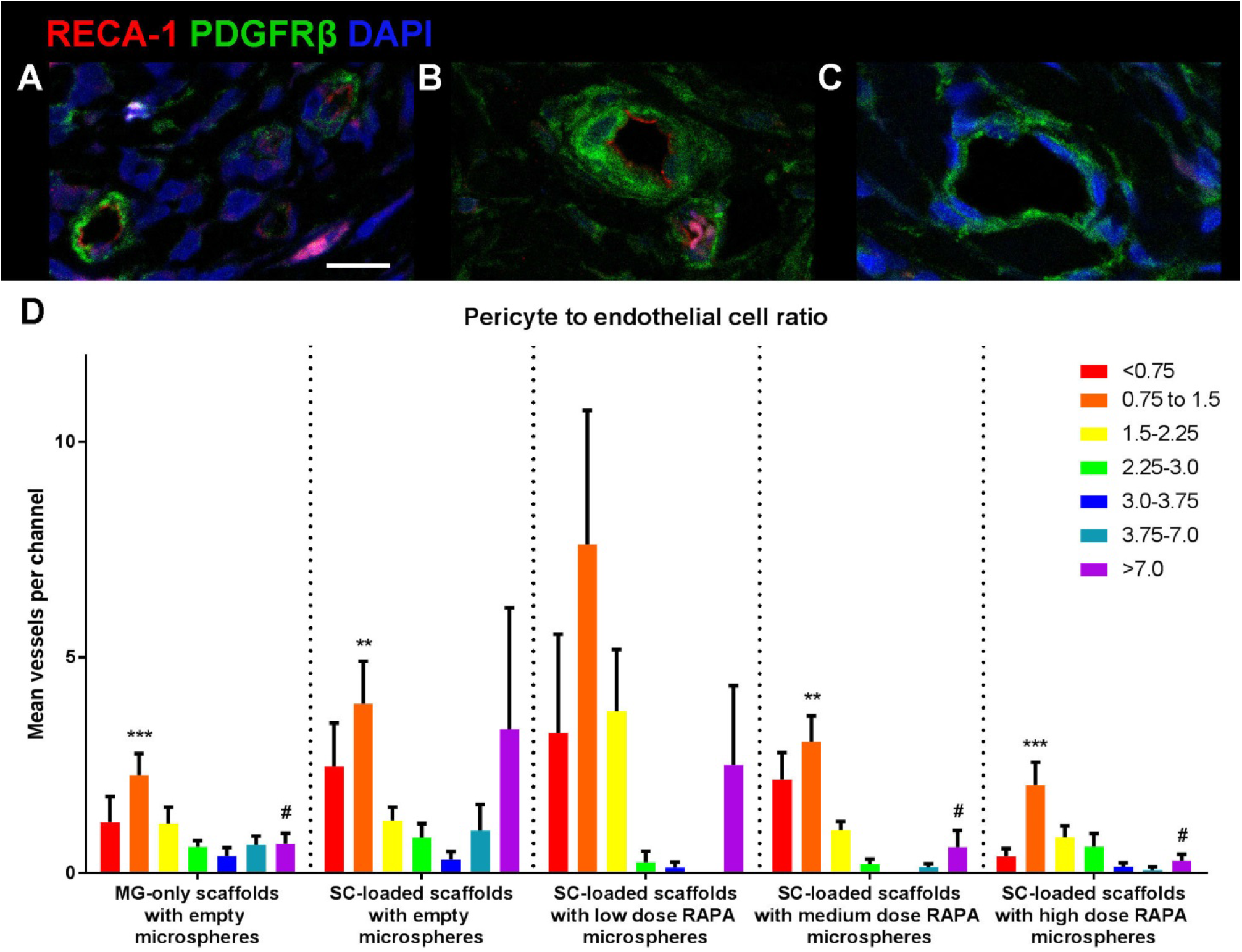
Vascular phenotype and function within scaffold channels *in vivo*. Representative high power confocal images of blood vessels stained for endothelial cells with RECA-1 (red) and pericytes with PDGFRβ (green). (A) Regular smaller diameter vessels with endothelial cells covered by pericytes characteristic of mature microvasculature in the channels of animals implanted with SC-loaded scaffolds with medium dose rapamycin microspheres. (B, C) Large irregular vessels with hypertrophic pericytes and vessel structures composed only of pericytes without endothelial cells were observed within SC-loaded scaffolds with empty microspheres. (A-C) 20 μm scale bar. (D) Ratio of pericyte/endothelial cell coverage of capillary lumina estimated by quantitative stereological measurement.

Pericyte to endothelial cell ratio (PC/EC ratio) was calculated for each vessel in a channel. Animals in the SC+Empty-MS group had the highest number of vessels with PC/EC >7.0, (Fig. 4 D). This immature vascular component was found to decrease with increasing RAPA concentration, and was also significantly lower in the MG+Empty-MS group. Animals in the SC+Low RAPA-MS group had the highest number of vessels with a PC/EC ratio of 0.75 to 1.5, representing a normal PC/EC ratio for healthy CNS vasculature (Fig. 4 D).

Vessel function was assessed in the second cohort of the study by tail vein injection of TRITC-dextran (Table 1). Foci of TRITC dye were rarely observed in the central channels of animals implanted with MG+Empty-MS (Fig. 5A). In contrast, the channels of animals implanted with SC+Medium RAPA-MS demonstrated many foci of concentrated TRITC dye as well as areas of more diffuse TRITC expression (Fig. 5B). Surface area measurements demonstrated a more than four-fold increase in TRITC dye signal in the channels of rapamycin treated animals (113.0 ± 32.04 μm^2^) compared with the Matrigel-only with empty microspheres group (26.34 ± 6.27 μm^2^) demonstrating improved vascular connectivity to the systemic circulation.

**Fig. 5.**
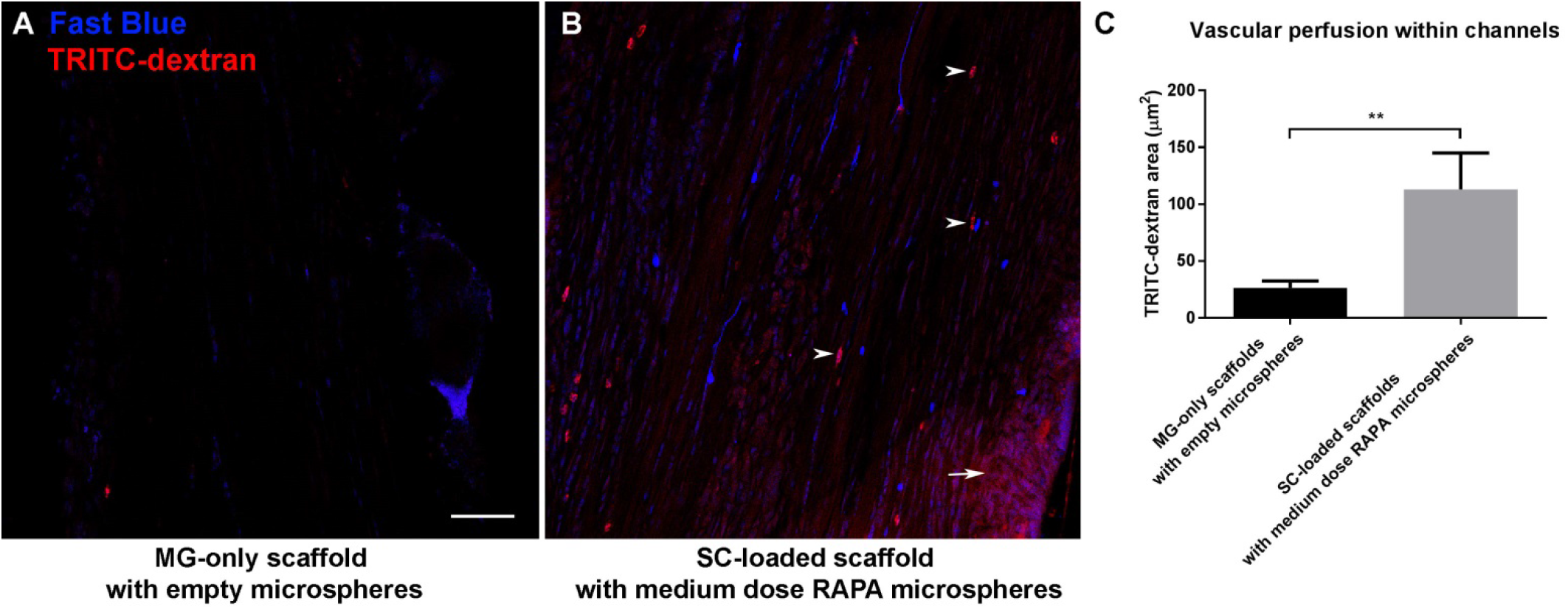
Representative confocal images of the innermost central scaffold channel in longitudinal sections. (A) Few foci of TRITC-dextran dye and Fast Blue labeled axons in Matrigel-only scaffolds with empty microspheres group. (B) Multiple TRITC-dextran dye foci in punctate, intravascular (arrow heads) and diffuse leakage patterns were observed in SC-loaded scaffolds with medium dose rapamycin microspheres along with longitudinally-oriented axons labeled with Fast Blue. 50 <m scale bar in panel (A) applies to panel (B). (C) Vascular perfusion quantitated using surface area measurements of intense TRITC dye (**p<0.01).

### Rapamycin treatment results in increased axonal regeneration across implanted scaffolds

One week after FB injection in animals in cohort 2, FB-labeled neuron nuclear profiles were counted within 15 mm spinal cord segments designated P1 – P3 rostral and S1 – S2 caudal (Fig. 6A, B). Cells with morphology resembling large motor neurons and smaller spinal interneurons were identified (Fig. 6 C, D). The Abercrombie formula correction factor was calculated as 0.802. The mean total of neurons rostral to the scaffold (P3 + P2 + rostral P1 segments) after caudal FB injection was higher in SC+Medium RAPA-MS group (36,326 +/-15,176 neurons, n=5 animals) compared to MG+Empty-MS animls (9,551 +/-1857 neurons, n=5 animals) (p=0.032) (Fig. 6E). The mean number of neurons counted in the caudal cord (caudal P1 + S1 + S2) after rostral dye injection was 5,602 +/-676 in the MG+Empty-MS group (n=5 animals), and 10,592 +/-2780 in the SC+Medium RAPA-MS group (n=4 animals) (p=0.413) (Fig. 6 F). The majority of counted neurons were in the rostral P3 and P2 segments in animals implanted SC+Medium RAPA-MS scaffolds, whereas the neuron counts in animals without implanted cells or rapamycin were more evenly distributed (Fig. 5G).

**Fig. 6.**
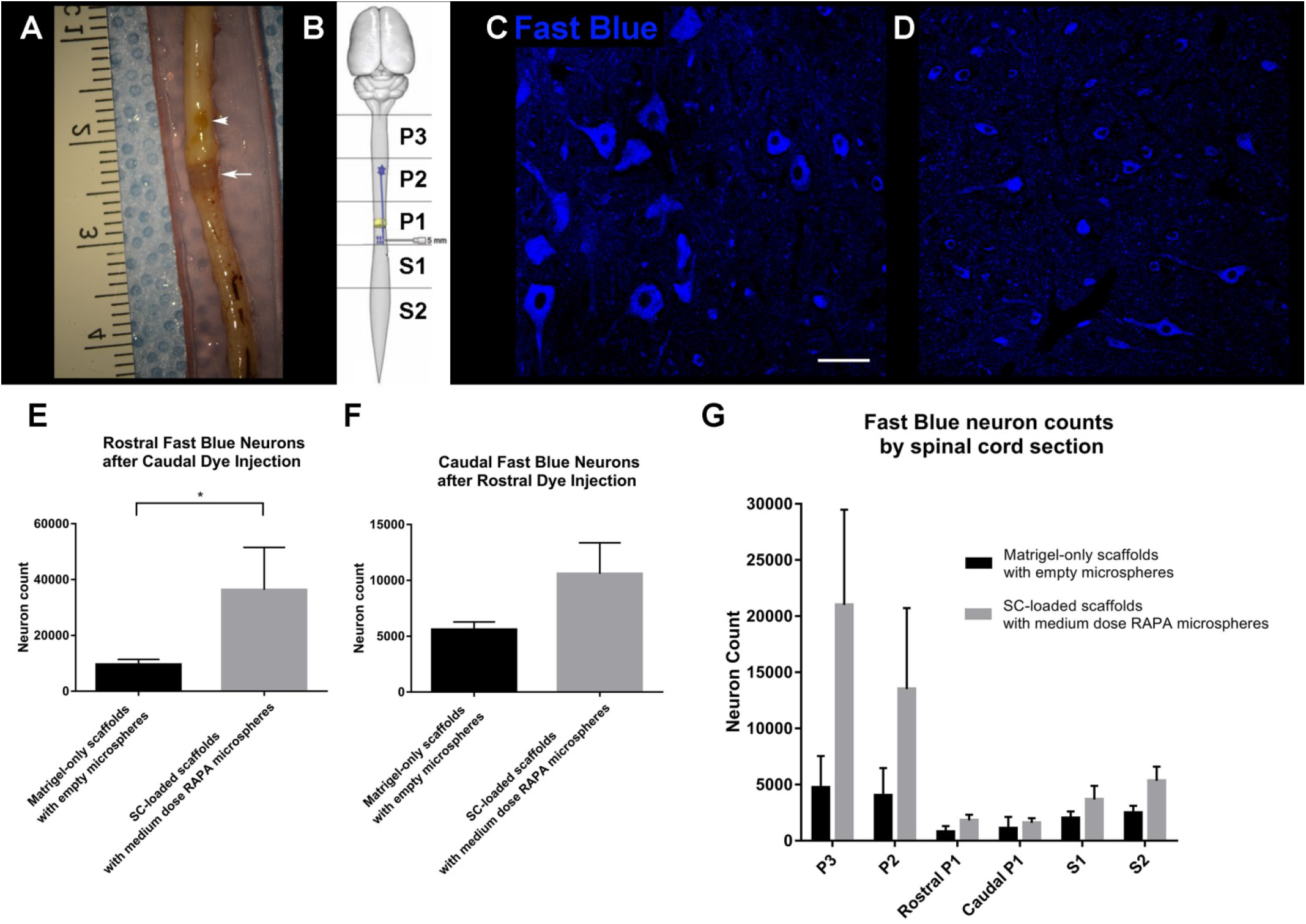
Retrograde tracing of regenerated axons with Fast Blue dye. (A) Gross image of the spinal cord of a scaffold-implanted animal with rostral injection of Fast blue dye. Tissue is oriented with rostral up and caudal down, with a metric ruler laid beside for scale. The site of Fast Blue injection can be seen as the tan spot (arrow head) rostral to the translucent OPF+ scaffold (arrow). (B) A schematic demonstrating the labeling of spinal cord segments. (C) Representative images of Fast Blue labeled neurons with characteristics predominately of large motor neurons and (D) characteristics predominately of small spinal interneurons. 50 μm scale bar in panel (C) applies to panel (D). (E) Quantification of total regenerated neurons labeled with Fast Blue in the combined rostral P1, P2 and P3 spinal cord segments (*p=0.05). (F) Combined caudal P1, S1, and S2 spinal cord segments following caudal and rostral Fast Blue injections, respectively. (G) Quantification of regenerated Fast Blue labeled neurons within each individually counted spinal cord segment (G).

## Discussion

Biomaterial based tissue engineering can bring together components of a system that have been individually demonstrated to have properties suitable for solving a complex problem. Combinatorial approaches to restore function after spinal cord injury have been advocated for a number of years (Olson 2013). Systematic testing of individual components to mechanistically predict and optimize their contribution to the final product represents a traditional engineering strategy. In this study two different biomaterials (PLGA and OPF+), rapamycin and Schwann cells were combined. Different types of biomimetic synthetic scaffolds have previously been compared and demonstrated that OPF+ supports the most robust axonal regeneration (Chen *et al*. 2011). Schwann cells produce the extracellular microenvironment that supports regeneration of CNS axons and promote regeneration in OPF+ scaffolds (Chen *et al*. 2017; Olson *et al*. 2009). Fibrosis and a foreign body response to SC-loaded OPF+ scaffolds were identified as a major remaining barrier to axon regeneration and functional recovery (Hakim *et al*. 2015). Rapamycin has been used to reduce fibrosis in reaction to various implanted synthetic materials (Morice *et al*. 2002; Moses *et al*. 2003).

We now demonstrate that combining SC with OPF+ scaffolds that contain PLGA microspheres providing sustained release of rapamycin restore function. This was associated with normalization of blood vessel formation and improved perfusion, increased axonal regeneration and reduced scar formation.

Initial studies to characterize and optimize sustained release of rapamycin from PLGA microspheres and demonstrate required biological activity for neurons, SC and fibroblasts were performed *in* vitro. Formulation of PLGA with a 50:50 ratio of lactic to glycolic acid demonstrated sustained release over 5 weeks. This was the target interval for the proposed short-term recovery studies *in vivo*. Previous work has demonstrated the reversible nature of fibroblast inhibition by rapamycin (Ong *et al*. 2007). The release kinetics of rapamycin from OPF+ scaffolds may be modified by the addition of PLGA microspheres formulated with different ratios of lactic to glycolic acid (Jain 2000). In future long-term experiments it will be necessary to characterize release kinetics of rapamycin delivery and the time course of the foreign body response to determine whether further improvements in locomotor function and axon regeneration are achieved with longer duration of rapamycin release.

Differences in SC morphology *in vivo* and neurotrophic factor production *in vitro* were observed with rapamycin treatment. Rapamycin has been previously shown to increase SC migration and nerve growth factor (NGF) secretion *in vitro* (Liu *et al*. 2014) In the current study, rapamycin treatment was also observed to increase BNDF and GDNF secretion by SC *in vitro*. The increases in BDNF and GDNF secretion by transplanted SC could contribute to the observed improvements in axon regeneration and functional recovery with rapamycin delivery from OPF+ scaffold *in vivo*. Further, the observed differences in SC morphology suggest that rapamycin may influence maturation of SC within the lesion environment. A recent study demonstrated that reduction in volume of SC cytoplasm that accompanies myelin maturation is regulated through autophagy and could be influence by rapamycin (Jang *et al*. 2015). The smaller, spindle shaped morphology of p75(NTR)-positive SC in rapamycin-containing scaffolds may suggest a more mature myelinating phenotype when compared to the larger, amoeboid morphology observed in scaffolds without rapamycin delivery.

The vascularization pattern within OPF+ scaffold channels was found to be dependent on whether scaffolds were loaded with SC and the dose of rapamycin delivered through PLGA microspheres. The observation that few vessels were present in the channels of MG-only scaffolds suggests that angiogenic factors produced by SC are crucial for implant vascularization. The neurotrophic factor BDNF, which is produced by SC *in vitro,* is known to be a potent inducer of angiogenesis (Kermani and Hempstead 2007). Of note, BDNF administration was able to stimulate *in vivo* neovascularization within Matrigel plugs transplanted subcutaneously in mice (Kermani *et al*. 2005). The neovascularization was found to be mediated through local tropomysin receptor kinase B (TrkB) activation by BDNF on endothelial cells as well as the recruitment of pro-angiogenic circulating hematopoietic cells. Rat brain derived endothelial cells were observed to express receptors for neurotrophic factors BDNF, NGF, TrkB and p75(NTR), respectively. BDNF activation of TrkB was found to increase endothelial cell survival, tube formation in three-dimensional collagen matrices, and vascular endothelial growth factor (VEGF) receptor expression through activation of PI3K/Akt dependent signaling pathways. In contrast, propeptide NGF (pro-NGF) activation of p75(NTR) signaling before cleavage by metalloproteinases was shown to increase endothelial cell apoptosis. The differing effects of BDNF and pro-NGF warrant future studies to evaluate their relative secretion by transplanted SC *in vivo,* including the influence of rapamycin on expression ratio between the factors and their proteolytic cleavage. In combination with these previously described effects of BDNF on endothelial cell function, the observation of increased SC within SC-loaded scaffolds at 6 weeks post implantation compared to MG-only scaffolds implicate BDNF as an important potential mediator of neovascularization within OPF+ scaffolds following SCI.

In addition to potentially modulating vascularization through influencing SC function, rapamycin may have a direct effect on endothelial cells participating in graft angiogenesis. Previous studies in malignant tumors have revealed the importance of a balance between local pro-and anti-angiogenic factors in the development of functional vasculature. Anti-angiogenic therapies, such as administration of anti-VEGF antibodies, have been shown to result in smaller diameter, less permeable, and less tortuous tumor vessels with increased perfusion and oxygen delivery in mice bearing vascularized tumor xenografts or allografts (Hansen-Algenstaedt *et al*. 2000; Yuan *et al*. 1996). An explanation of the paradoxical improvements in vascular function with anti-angiogenic therapies has been the vascular normalization hypothesis. The hypothesis proposes that the overabundance of pro-angiogenic factors such as VEGF, TGFβ, and PDGF within the tumor microenvironment produce an imbalance in pro-and anti-angiogenic signaling leading to the development of abnormal microvasculature (Jain 2001; Jain 2005). Thus, more mature and functional vascular phenotypes are restored by exogenous anti-angiogenic treatments. Rapamycin has been shown to have antiangiogenic properties through reducing VEGF secretion by tumors and preventing VEGFR signaling in endothelial cells (Guba *et al*. 2002). Macrophages and foreign body giant cells in contact with biomaterials may overproduce angiogenic factors such as TGFβ and PDGF (Rappolee *et al*. 1988; Song *et al*. 2000). It is possible that a process of vascular normalization akin to that described in the cancer literature has occurred with rapamycin treatment in the current study.

This possible explanation of the improvements in vascularizaton observed with rapamycin treatment is supported by the similarities in perivascular cell phenotypes and vessel morphologies between tumor vasculature and that seen in channels of SC-loaded scaffolds with empty microspheres. Hyperplasia of pericytes is a defining characteristic of the microvasculature observed in high grade malignant gliomas (Sun *et al*. 2014). Vessel-like structures consisting only of cells with pericyte markers have also been observed in glioma tissue. SC-loaded scaffolds with empty microspheres exhibited similar findings suggesting that alternative mechanisms of pericyte-mediated vessel extension may be involved in the unregulated angiogenesis observed in both tumor and tissue engineering contexts. Comparable pericyte abnormalities to those observed in the OPF+ scaffolds have also been described in pancreatic, breast, and lung carcinomas (Morikawa *et al*. 2002).

Large diameter, irregularly shaped, and hyperpermeable vessels with abnormal pericyte coverage been produced in non-tumor tissue in transgenic mice with inducible expression of a myristoylated form of Akt1 (myrAkt1) (Phung *et al*. 2006). These large and irregular vessels were similar in appearance to those observed in SC-loaded OPF+ scaffolds with empty microspheres. The abnormal vessel phenotype formed over a period of 6-7 weeks upon myrAkt1 induction and reversed after myrAkt1 suppression for 4 weeks. Concurrent rapamycin treatment almost entirely prevented the development of abnormal vasculature following myrAkt1 induction. These results, may explain the similar effects in vascular normalization observed within OPF+ scaffolds when rapamycin is delivered from PLGA microspheres. Future studies evaluating Akt activity within the scaffold channels and mTOR dependent pathways will be important (Sarbassov *et al*. 2006). Identification of these mechanisms will provide important information that could lead to further improvements of vascularized biomaterial implants for CNS regenerative repair.

In conclusion, this study demonstrates the potential for tissue engineering to bring multiple regenerative strategies to restore function after spinal cord injury. This approach also allows multiple components of the system to be modulated in a controlled and independent way. This includes the site and rate of drug delivery. Microsphere composition can be manipulated to produce timed release at different periods and for different intervals. In the future different types of cells or cells releasing different growth factors in different regions of the scaffolds may be used to regionally regulate regeneration of specific classes of axons including ascending sensory or descending motor projections. Rational and mechanism-based approaches are available to systematically modulate these processes using combinatorial tissue engineering strategies.

## Acknowledgements

This work was supported by grants from the National Institute of Biomedical Imaging and Bioengineering (grant number EB02390); National Institute of Biomedical Imaging and Bioengineering (grant number 2T32EB005583); National Center for Advancing Translational Sciences (grant number TL1 TR002380); Morton Cure Paralysis Fund; and Craig H. Neilsen Foundation. The authors wish to acknowledge the expert technical assistance of Jarred Nesbitt and the expert administrative assistance of Jane Meyer.

---

**Fig. S1.**
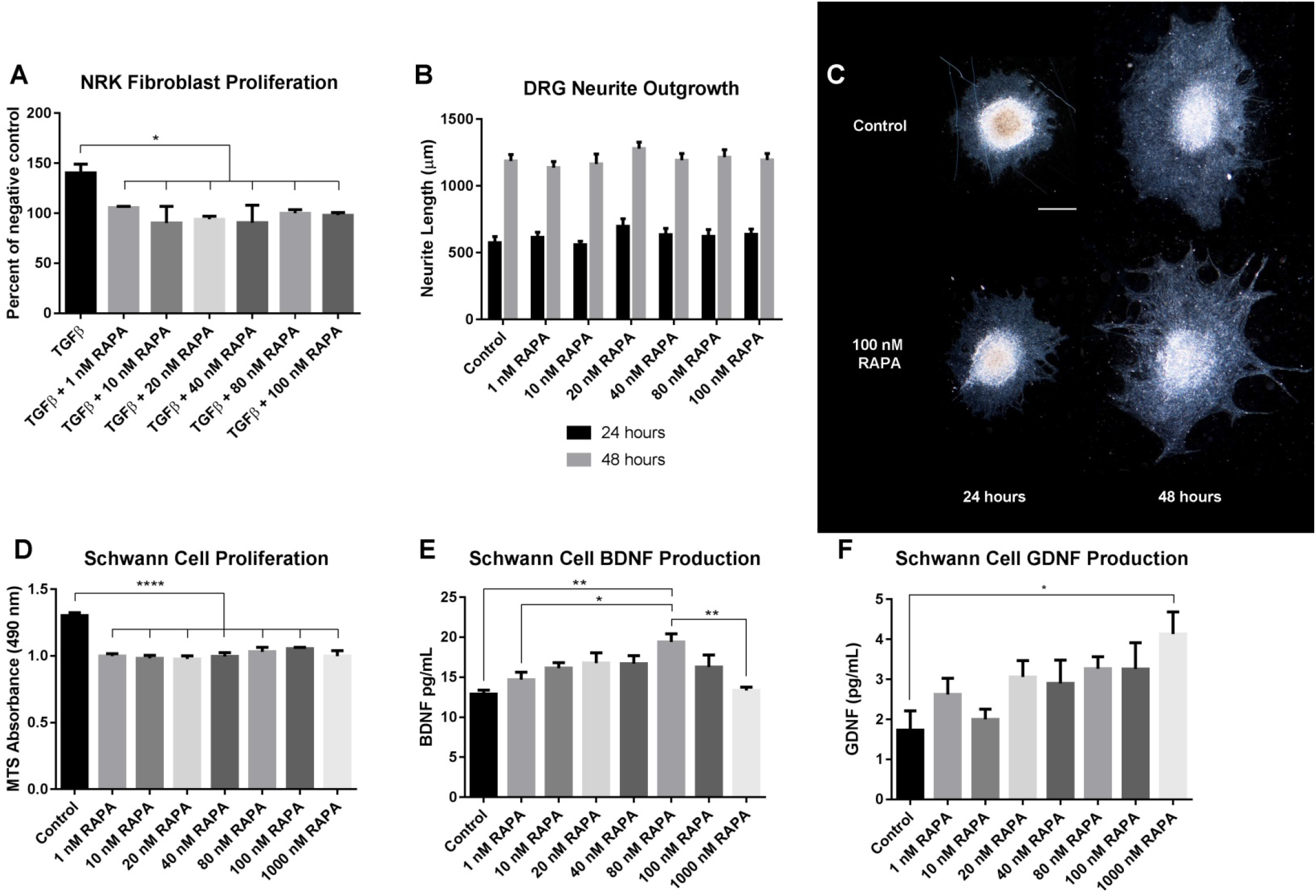
*In vitro* effects of rapamycin on fibroblasts, Schwann cells, and DRG Neurons. The effects of free drug rapamycin (A) and rapamycin-containing PLGA microspheres (B) in preventing TGFβ1-stimulated NRK fibroblast proliferation assessed by MTS assay *in vitro*. Proliferation as determined by absorbance at 490 nm is expressed as percentage of unstimulated serum starved negative control cells. (*) denotes significant difference from positive control TGFβ or TGFβ and empty microspheres groups in (A) and (B), respectively. (C) DRG neurite outgrowth in the presence of free drug rapamycin at 24 and 48 hours as assessed by longest neurite length. (D) Schwann cell proliferation in the presence of free drug rapamycin as determined by absorbance at 490 nm following MTS assay. (*) denotes significant difference from all rapamycin treated groups. Schwann cell production of BDNF (E) and GDNF (F) as determined by ELISA of Schwann cell conditioned media. In panel (E), (*) denotes significant difference from 80nM rapamycin group.

**Fig. S2.**
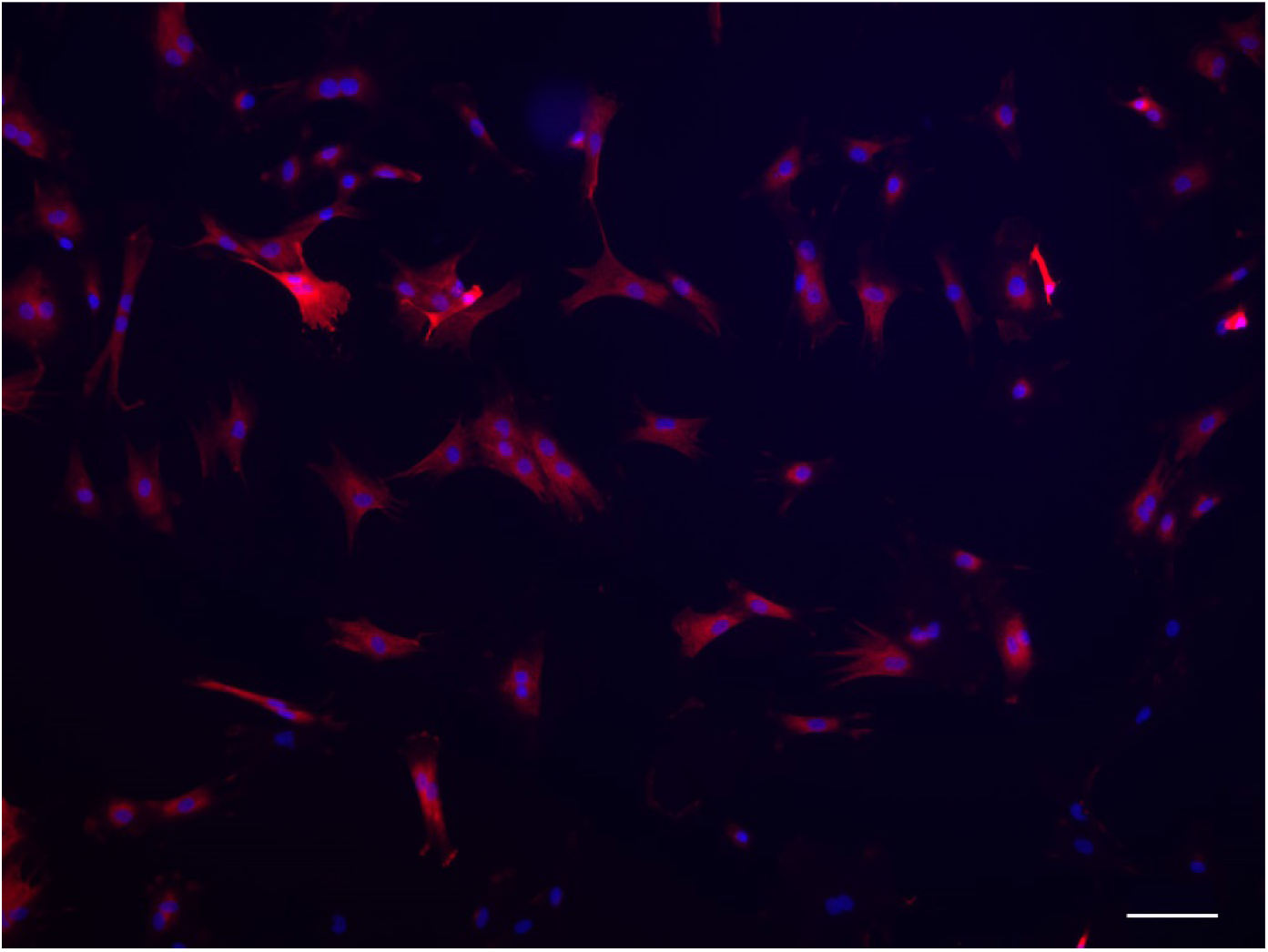
Representative image of p75(NTR) (red) expression in cultured neonatal rat primary Schwann cells after 4 passages *in vitro*. Cultures were counterstained with DAPI. The proportion of Schwann cells positive for p75(NTR) expression (phenotypic purity) was counted as a percentage of the total number of DAPI positive cells in 15 random fields.

**Fig. S3.**
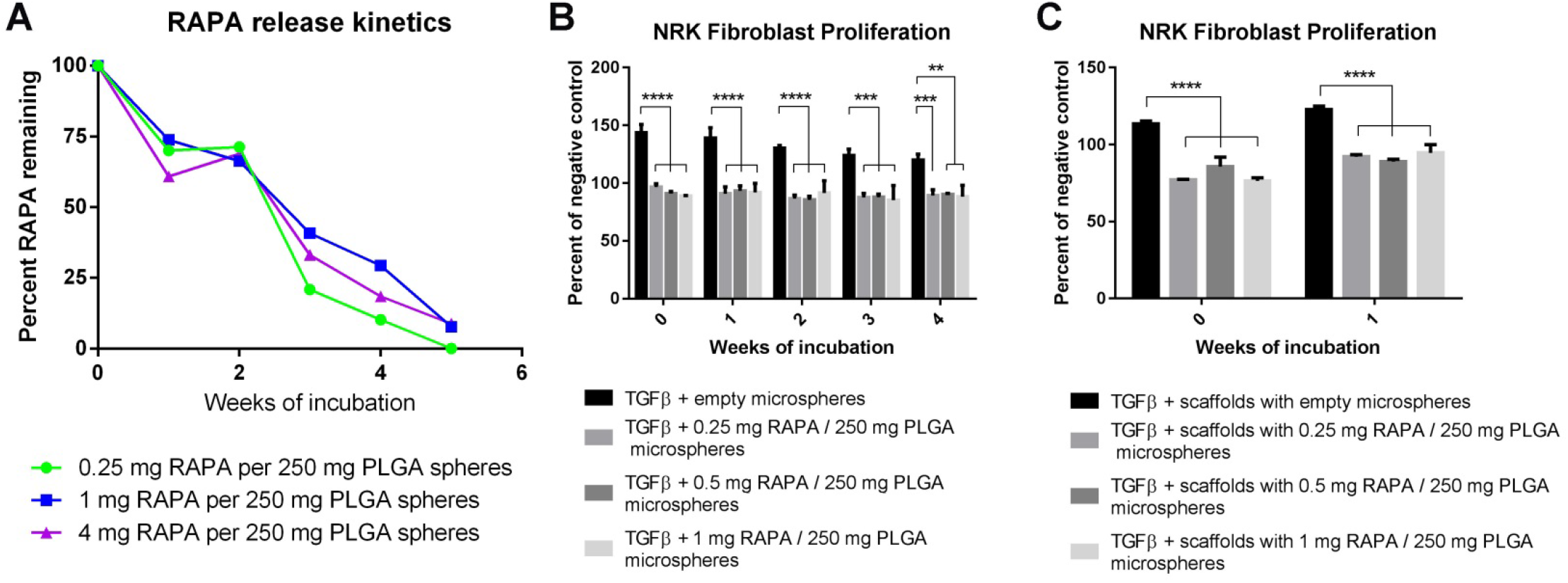
Rapamycin release kinetics from PLGA microspheres. demonstrate a burst release during the first week, a lag in release over the second week, and a steady release over weeks 3 through 5 (A). OPF+ scaffolds incorporating rapamycin-containing PLGA microspheres are able to prevent TGFβ1-stimulated NRK fibroblast proliferation *in vitro* as assessed by MTS assay (B). Proliferation as determined by absorbance at 490 nm is expressed as percentage of unstimulated serum starved negative control cells. (*) denotes significant difference from positive control TGFβ and scaffolds with empty microspheres groups.

**Fig. S4.**
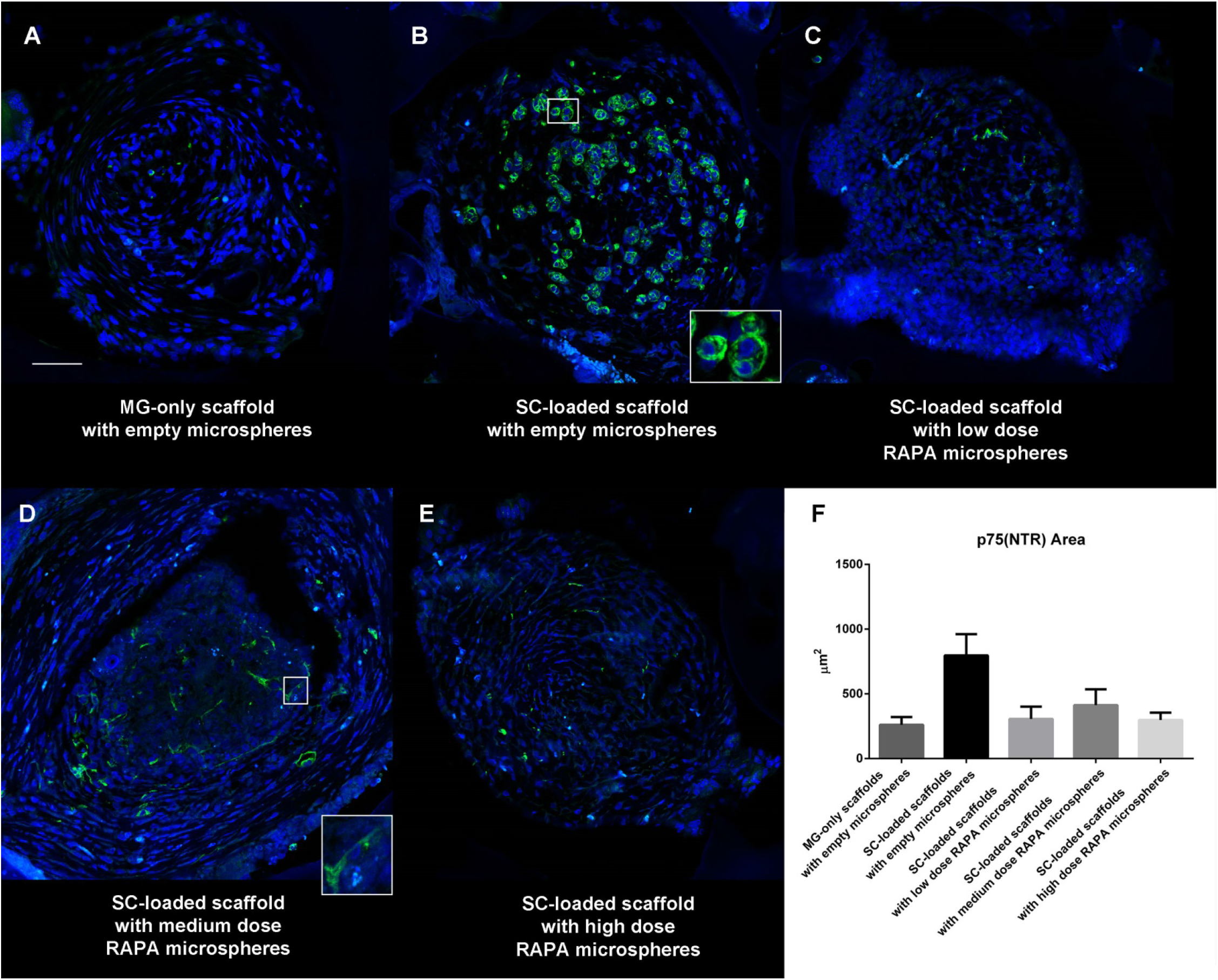
Confocal images of individual scaffold channels stained for Schwann cells with p75(NTR) demonstrate fewer Schwann cells at 6 weeks post implantation within Matrigel-only scaffolds with empty microspheres (A), when compared to the many amoeboid shaped Schwann cells within SC-loaded scaffolds with empty microspheres (B). Fewer Schwann cells are also observed in rapamycin treated animal groups (panels C though E), but these cells have a distinct spindle shaped morphology. 50 urn scale bar in panel (A) applies to panels (B) through (E). Quantification of p75(NTR) area within scaffold channels (F).

**Fig. S5.**
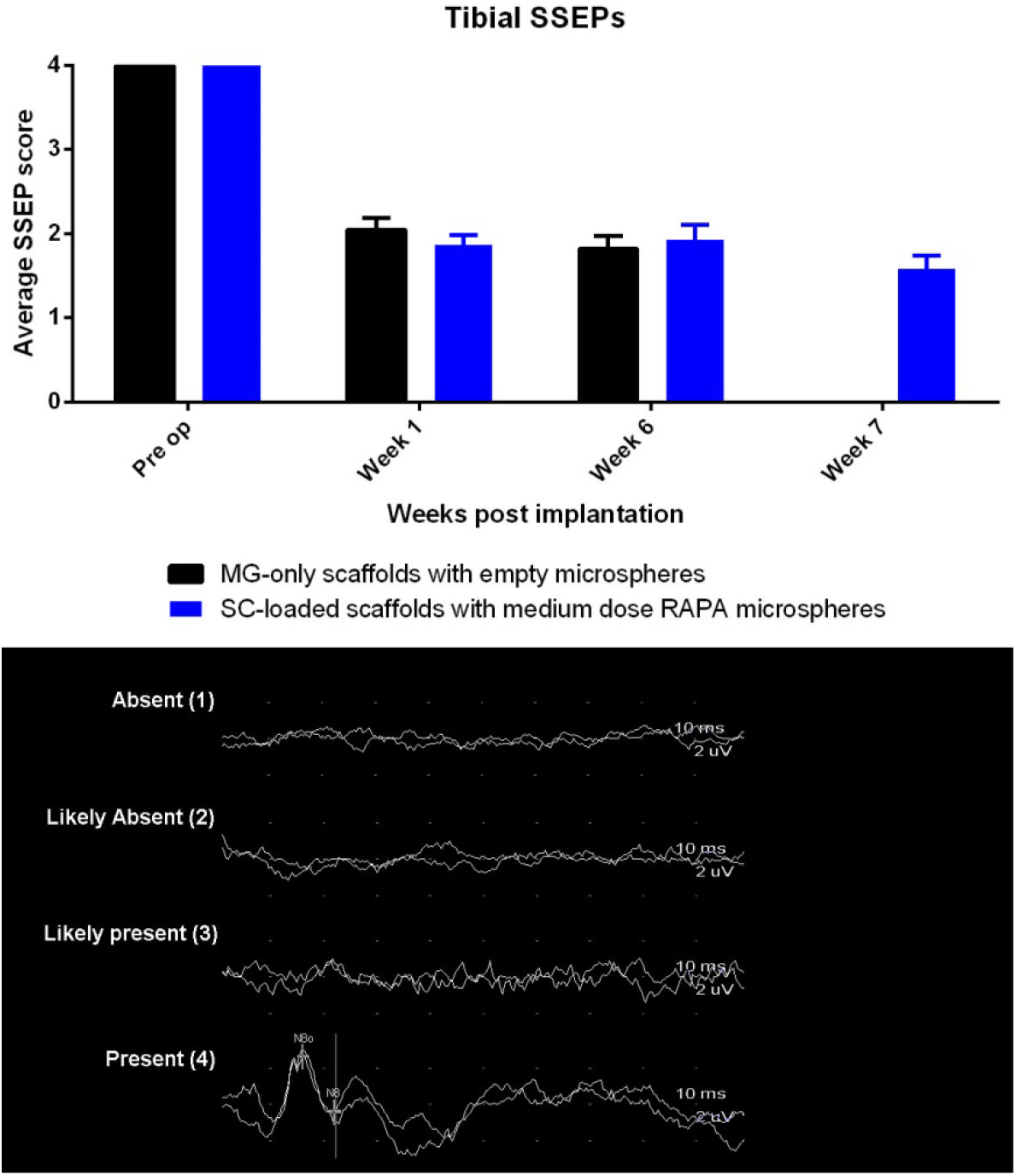
Averaged SSEP scores of animals from the second cohort. demonstrate a loss of tibial SSEPs following transection spinal cord injury and scaffold implantation. No significant recovery of SSEP waveforms were observed throughout the duration of the experiment. Representative waveforms demonstrating each of the numerical scores are shown.

